# AmpliDiff: An Optimized Amplicon Sequencing Approach to Estimating Lineage Abundances in Viral Metagenomes

**DOI:** 10.1101/2023.07.22.550164

**Authors:** Jasper van Bemmelen, Davida S. Smyth, Jasmijn A. Baaijens

## Abstract

Metagenomic profiling algorithms commonly rely on genomic differences between lineages, strains, or species to infer the relative abundances of sequences present in a sample. This observation plays an important role in the analysis of diverse microbial communities, where targeted sequencing of 16S and 18S *ribosomal RNA* (rRNA), both well-known hypervariable genomic regions, have led to insights into microbial diversity and the discovery of novel organisms. However, the variable nature of discriminatory regions can also act as a double-edged sword, as the sought-after variability can make it difficult to design primers for their amplification through *Polymerase Chain Reaction* (*PCR*). Moreover, the most variable regions are not necessarily the most informative regions for the purpose of differentiation; one should focus on regions that maximize the number of lineages that can be distinguished. Here we present AmpliDiff, a computational tool that simultaneously finds such highly discriminatory genomic regions, as well as primers allowing for the amplification of these regions. We show that regions and primers found by AmpliDiff can be used to accurately estimate relative abundances of SARS-CoV-2 lineages, for example in wastewater sequencing data. We obtain mean absolute prediction errors that are comparable with using whole genome information to estimate relative abundances. Furthermore, our results show that AmpliDiff is robust against incomplete input data and that primers designed by AmpliDiff continue to bind to genomes originating from months after the primers were selected. With AmpliDiff we provide an effective and efficient alternative to whole genome sequencing for estimating lineage abundances in viral metagenomes.

## Background

Studying the composition of metagenomic samples through genome sequencing has led to insights into diverse microbial communities [46, 11, 42]. While whole (meta)genome sequencing is frequently used for this purpose, not all regions of the genome are equally informative. If the goal is to distinguish between different lineages, strains, variants, or species, we should focus on those regions that allow us to discriminate between the genomes of interest. For example, many studies focus only on 16S (bacteria and archaea) or 18S regions (eukaryotes) for metagenomic profiling[39, 38]. Such a targeted sequencing approach has two main advantages: first, it is cost-effective as it requires fewer sequencing reagents per sample, and second, it reduces the computational burden in downstream processing due to generating less data that needs to be processed.

Targeted sequencing, otherwise known as amplicon sequencing, is a method in which genetic material of interest in (meta)genomic samples is amplified, using *Polymerase Chain Reaction* (*PCR*) for example, and later sequenced. Depending on the application, the objective can be to identify the presence of particular genetic markers [13], or to differentiate between different species that are present [52, 59] for example. Moreover, if one is interested in *Whole Genome Sequencing* (*WGS*) it is also possible to design primers such that the resulting PCR products span the entire genome(s) of interest.

Amplicon sequencing is widely applied in the context of *Wastewater-Based Epidemiology* (*WBE*) [1, 24, 37, 51, 57]. In wastewater-based epidemiology, the objective is to study population-scale phenomena, such as trends in the number of infections based on corresponding pathogen concentrations found in wastewater samples [10]. Since this method is non-invasive and captures information from the entire population within a geographical area, it is less prone to biases that occur in clinically obtained samples [3]. However, wastewater samples can suffer from low concentrations of genetic material, environmental RNA degradation, the presence of inhibitors, and high fragmentation rates [43]. For this reason, amplification of the genetic material of interest is a crucial step in wastewater analysis.

The importance of WBE itself has become apparent during the SARS-CoV-2 pandemic. A multitude of studies have confirmed that SARS-CoV-2 can be detected in wastewater samples [5] and that we can estimate COVID-19 case numbers from such samples [33, 40, 41, 60]. Moreover, we can estimate the relative abundances of different SARS-CoV-2 lineages (and therefore different variants) from wastewater samples using amplicon-based sequencing [4, 15, 28, 30, 47, 48, 63], further emphasizing the applicability of amplicon-based sequencing. The key step in performing amplicon-based sequencing is designing a set of primers such that the genetic material of interest can be amplified [14]. Several physicochemical constraints (e.g. melting temperature, inter-primer interaction, intra-primer interaction) have to be considered, for which several different primer design tools (e.g. Primer3 [54], Primer-BLAST [62], PriMux [22] and openPrimeR [34]) have been developed that take all constraints into account. These tools take as input a set of reference genomes and a region of interest and attempt to find a set of primers such that the region of interest can be amplified in as many reference genomes as possible. In addition, PrimalScheme [44] and Olivar [58] have the option to find a near-minimal size set of primers such that the resulting amplicons would cover the entire genome, enabling amplicon-based WGS.

These primer design tools have been successfully applied in practice, but they can only find primers once a region of interest has been provided, or when the goal is to amplify the whole genome. This suffices when such regions are known a priori, for example in 16S sequencing [39] and 18S sequencing [18]. However, when this is not the case there is a need for tools that can find regions of interest, along with primers. Applications can include the detection of pathogens [13], or facilitating the estimation of relative abundances of species present in a sample by focusing on variable regions in the genomes [50]. In the latter case, the focus should be on finding amplicons and corresponding primers such that the resulting amplicons can be used to differentiate between species in a sample. These steps should be considered simultaneously, as potential amplicons are only useable if we can find feasible primers in the flanking regions; effectively, the problem of determining a discriminatory set of amplicons depends on the primer finding problem. To our knowledge, there is no tool that integrates finding amplicons and corresponding primers for optimal differentiation between different genomic classes.

Here we introduce a new methodology, AmpliDiff, that identifies discriminatory regions in a set of genomes and simultaneously finds primers such that these regions can be amplified through PCR. By considering multiple input genomes, AmpliDiff can find primers in relatively conserved regions, such that the resulting PCR products are able to discriminate between different genomes (or classes thereof). We demonstrate AmpliDiff by computing amplicons and corresponding primers for SARS-CoV-2 genomes. The effectiveness of these amplicons is assessed by evaluating the abundance estimation accuracy for different SARS-CoV-2 lineages on simulated data, comparing estimates on only selected amplicons versus whole genome sequencing. Finally, by showing that these primers also match genomes discovered months after the primers were generated, we conclude that AmpliDiff provides a robust, efficient, and economic alternative to whole genome amplification that can be applied in the context of metagenomic profiling.

## Results

### An overview of AmpliDiff

AmpliDiff is designed to find a minimal set of amplicons, along with corresponding primers, to differentiate between classes in a given set of input genomes. The algorithm consists of three steps: (i) extracting a set of candidate primers, (ii) finding candidate amplicons and determining their ability to discriminate, (iii) greedily selecting amplicons and validating whether it is possible to find a set of primers for these amplicons. Figure 1 outlines the overall methodology of AmpliDiff which we will briefly describe here, with a more thorough explanation available in the Methods section.

**Figure 1:**
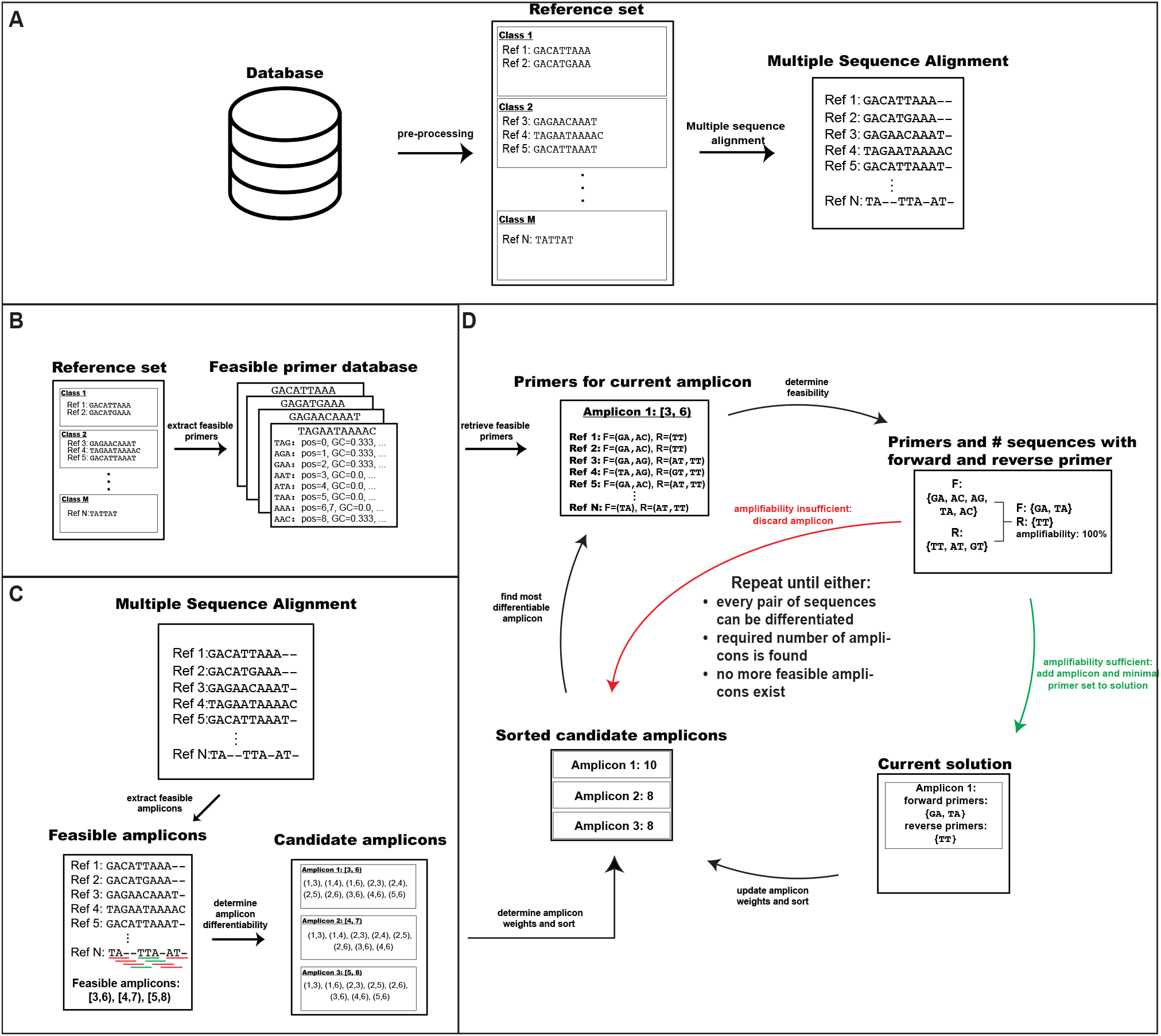
Outline of the workflow of AmpliDiff. **(A)** Pre-processing consists of building a reference set from a database of genomes (e.g. GISAID) and applying multiple sequence alignment to obtain multiple aligned genomes. **(B)** AmpliDiff builds a database of feasible primers that can be considered during the greedy amplicon selection (iii) step. **(C)** Feasible amplicon candidates are extracted from the multiple aligned genomes, and for every amplicon, the pairs of genomes it can differentiate are stored. **(D)** In its final step, AmpliDiff greedily finds the most discriminatory amplicon, retrieves flanking primers, and determines if a (sub)set of primers can be found to amplify the amplicon in the reference genomes. If this is possible, the set of primers is minimized and the amplicon and primers are added to the solution, after which the discriminatory power of amplicons is updated and the amplicons are sorted accordingly. If amplification is not possible, the amplicon is discarded and the next most discriminatory amplicon is considered. This process is repeated until either all pairs of genomes can be differentiated, a user-defined number of amplicons is found, or no more feasible amplicons exist. Finally, AmpliDiff outputs the selected amplicons and corresponding primer sets.

The input to AmpliDiff consists of multiple aligned genomes in the form of a *Multiple Sequence Alignment* (*MSA*), each with a corresponding class. AmpliDiff then first extracts every physicochemically feasible primer of a given length and stores them in a *primer database* (Figure 1**B**) to be used in step (iii). Next, candidate amplicons are found and their differentiability is determined (Figure 1**C**). We define the *differentiability* of an amplicon as the number of pairs of genomes of different classes that it can differentiate. In its final step, AmpliDiff heuristically tries to find a minimal set of amplicons with corresponding primers to differentiate between all the pairs of genomes. This is done by iteratively finding the most differentiable amplicon and checking if it is possible to find forward and reverse primers to amplify it in the input genomes (Figure 1**D**), given the physicochemical constraints (see Methods). Defining the *amplifiability* of an amplicon as the relative number of genomes in which both a forward and reverse primer flanking the amplicon can be found, AmpliDiff only accepts amplicons that meet a user-provided minimal required amplifiability (default = 95%). If this amplifiability is met, AmpliDiff minimizes the number of primer pairs needed to amplify the amplicon and adds the amplicon along with a minimal set of primers to the solution. Afterward, the differentiability of the remaining candidate amplicons is updated to correct for the pairs of genomes that can already be differentiated, and the amplicons are sorted (highest to lowest) based on the newly calculated differentiabilities. When the amplifiability of an amplicon is too small, the amplicon is discarded and the next most differentiable amplicon is considered. This cycle is repeated until either every pair of genomes can be differentiated, a predefined number of amplicons has been found, or there are no more candidate amplicons. The final output of AmpliDiff consists of the resulting set of amplicons (with coordinates based on the provided MSA) along with the corresponding forward and reverse primers.

### AmpliDiff finds highly differentiable amplicons in SARS-CoV-2 genomes

Using AmpliDiff we have determined two sets of 10 amplicons: a set of 200 *basepair* (*bp*) wide amplicons, and a set of 400 bp wide amplicons (corresponding to Illumina iSeq 100 and MiSeq sequencing respectively). As input, we used a global reference set of 2749 SARS-CoV-2 genomes over all 1837 different lineages (1-7 genomes per lineage) that existed at the time that these sequences were downloaded (18 August 2022). The reference set was constructed by following the pre-processing steps described in the VLQ pipeline [4], using all of the available genomes on GISAID [31], and were multiple aligned with MAFFT [29]. Every genome on GISAID comes with an assigned Pango lineage [45]. These lineages consist of closely related SARS-CoV-2 genomes which are defined by key phylogenetic markers and shared mutations. Our objective is to find amplicons that allow us to differentiate between these lineages.

Figure 2**(a)** shows the relative differentiability of all amplicons of a fixed width of 200 and 400 bp, respectively, with at least 50 bp flanking on either side. Clearly, any 200 bp amplicon contained by a 400 bp amplicon, is at most as differentiable as the 400 bp amplicon. We observe that both the open reading frame 1a (ORF1a) and ORF1b regions are relatively conserved between the reference genomes and that regions close to the 3’-end (right) of the genome are generally more differentiable than regions closer to the 5’-end (left), with amplicons reaching up to 93.3% differentiability. It is possible that different amplicons can differentiate between the same pairs of genomes, and as such the figure does not tell us anything about which amplicons to combine in order to attain a shared differentiability of 100%. Moreover, it is not guaranteed that every amplicon is practically feasible since we need primers flanking the amplicons in order to amplify these regions through PCR. This can be difficult, as highly differentiable amplicons are generally located in relatively variable regions of the genome. Hence the width of the variable region can become a limiting factor: a width larger than the desired amplicon might prevent us from finding feasible primers flanking the amplicon.

**Figure 2:**
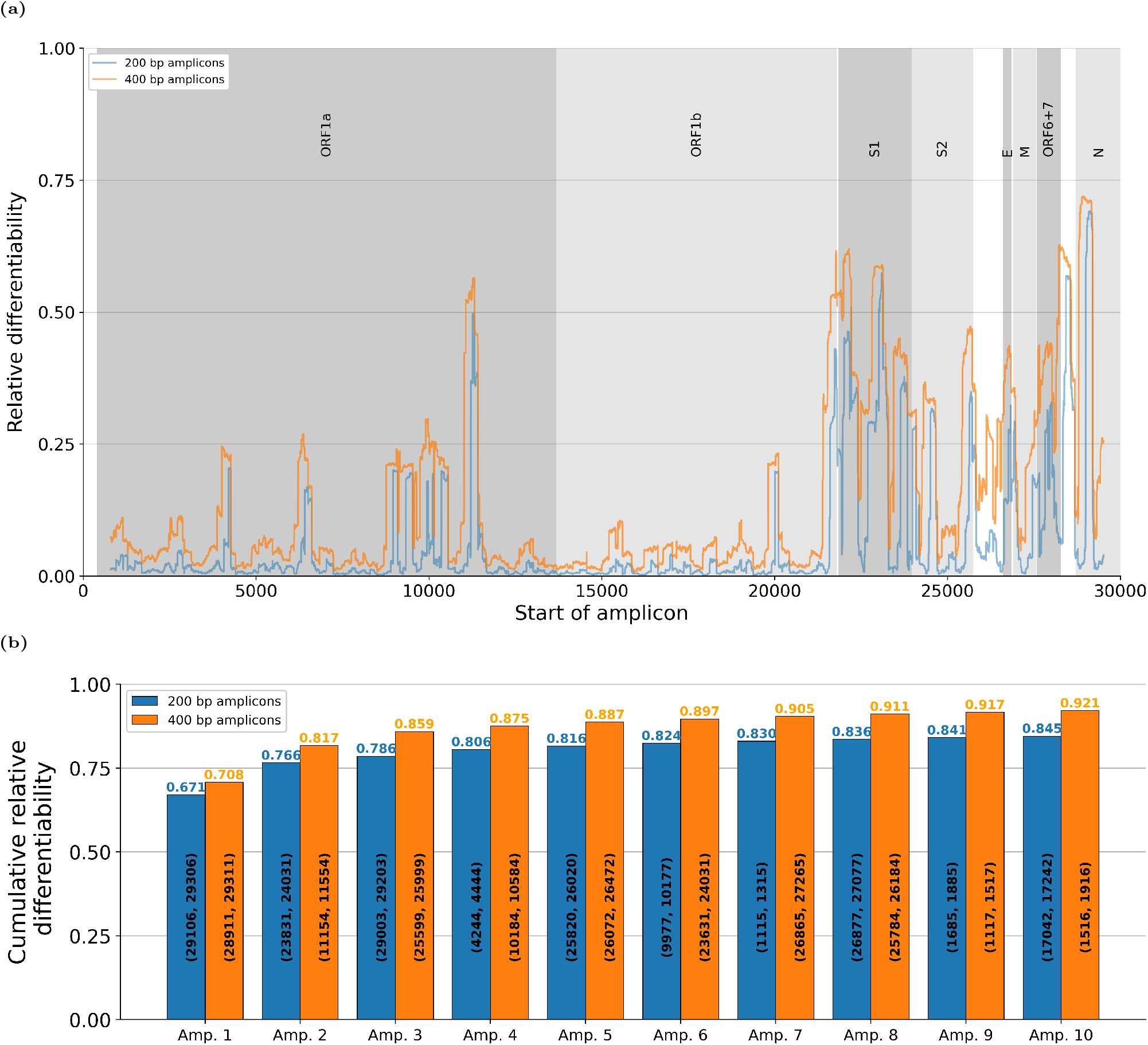
Overview of the differentiability of potential amplicons, and the amplicons selected by AmpliDiff on the global SARS-CoV-2 dataset. **(a)** Relative differentiability of all amplicons of widths 200 bp (blue) and 400 bp (orange) that have at least 50 nucleotides before or after them. **(b)** Cumulative relative differentiability of the ten best amplicons (200 and 400 bp, respectively) found by AmpliDiff.

The amplicons selected by AmpliDiff (10 amplicons with fixed widths of 200 and 400 bp, respectively, and a minimal amplifiability of 95%) are shown in Figure 2**(b)**. For both amplicon widths, we see that the first amplicon is chosen in the gene that encodes the nucleocapsid protein (N), which provides a relative differentiability of 67.1% and 70.8% for amplicons of 200 and 400 bp, respectively. Subsequent amplicons are spread across the genome, but in both cases, we observe that at least one amplicon in the spike gene (S)—a region with high differentiability—was selected. In the context of wastewater sequencing, this region often suffers from low sequencing depths (amplicon drop-out) which makes it a difficult target in practice [6]. This may be due to primer binding issues: in Figure 2**(a)**, we observe that the variation between different SARS-CoV-2 lineages in the spike gene is spread over a relatively wide region, making it difficult to find primers that bind to all sequences, even though the amplicon would be highly differentiable.

Another interesting observation is that we find some overlapping amplicons. While this may seem counter-intuitive, it can be the result of our primer minimization model which includes a trade-off between the number of primers required, and the number of genomes in which the amplicon can be amplified (i.e. has both a binding forward and reverse primer). This can lead to AmpliDiff suggesting overlapping amplicons in order to differentiate between the sequences for which one amplicon was not amplifiable due to requiring too many primer pairs. Alternatively, it is possible that variability is contained in a region that is wider than the pre-defined amplicon width, in which case at least 2 overlapping amplicons can be required to capture it. This latter possibility appears to be the case here, as even when requiring an amplifiability of 100%, overlapping amplicons are found (Figure 6 in the Supplementary Material).

We also see that for both the 200 and 400 bp amplicons, the added differentiability seems to drop exponentially with every additional amplicon. Conceptually this makes sense, as we expect every subsequent amplicon to be less differentiable than the previous—this is how the greedy algorithm picks amplicons. As a consequence, we see that the first 5 amplicons already allow us to differentiate between 81.6% and 88.7% of the pairs of genomes of different lineage for widths 200 and 400 bp, respectively. Any further amplicons contribute approximately 0.4-1% to the cumulative relative differentiability indicating that, depending on the required resolution, it can be sufficient to use up to 5 amplicons.

#### The effect of different minimal required amplifiabilities

The amplicons considered so far are based on a minimal required amplifiability of 95%. To get an idea of how the minimal amplifiability affects the amplicons selected, and in particular the number of primers required, we also ran AmpliDiff with minimal required amplifiabilities of 90%, 92.5%, 97.5%, 99.9% and 100%. The resulting cumulative differentiabilities for every respective configuration can be found in Figures 1-6 in the Supplementary Material. For minimal amplifiabilities of below 99.9% we see that roughly the same regions were selected, whereas for higher amplifiability thresholds the strictness in requirements becomes a bottleneck, and different amplicons are chosen. Nevertheless, we see that the initial amplicon is always located in the nucleocapsid gene and that the first few amplicons contribute the most to the cumulative differentiability.

An interesting observation that can be seen in both the 99.9% and 100% minimal amplifiability results, is that the first amplicon has a larger relative differentiability than the first amplicons for the lower amplifiabilities. This is due to the fact that our primer minimization model includes a trade-off between the number of primer pairs selected and the additional differentiability gained by adding them. Even though any solution allowed for higher amplifiability requirements is also valid when the required amplifiability is lower, the model can decide to not add primer pairs if they allow for only a marginal increase in differentiability. Hence, the same amplicon can achieve slightly lower differentiability depending on the primers selected.

The impact of requiring a higher minimal amplifiability on the number of primer pairs can be seen in Table 1 in the Supplementary Material where, for every configuration, we show the minimal number of forward and reverse primers required to amplify the amplicon found by AmpliDiff. Results are mostly similar for amplifiabilities of below 97.5%, where generally only a single primer pair is needed for the amplification of an amplicon in the corresponding number of input genomes. For minimal required amplifiabilities of at least 97.5%, however, we see that the number of required forward and reverse primers increases substantially, requiring up to 13 forward primers or 9 reverse primers when the minimal required amplifiability is 100%.

**Table 1:** Mean absolute prediction errors for both the amplicon-based approach (1, 2, 5, 10 amplicons) and the whole genome-based approach using fragment lengths of 200 and 400 over 20 simulations in both datasets when lineages are cut-off and aggregated at most 3 sublineage levels. In parentheses, the standard deviations are given.

| Amplicons | 200 bp amplicon width |  |  |  |  | 400 bp amplicon width |  |  |  |  |
| --- | --- | --- | --- | --- | --- | --- | --- | --- | --- | --- |
|  | 1 | 2 | 5 | 10 | WGS | 1 | 2 | 5 | 10 | WGS |
| MAPE (TX) | 3.619 (0.001) | 1.391 (0.001) | 2.729 (0.004) | 0.638 (0.006) | 3.674 (0.020) | 5.049 (0.001) | 3.842 (0.017) | 1.960 (0.021) | 0.936 (0.008) | 3.718 (0.022) |
| MAPE (NL) | 2.468 (0.005) | 5.757 (0.002) | 2.277 (0.008) | 0.760 (0.011) | 0.845 (0.028) | 1.060 (0.053) | 1.476 (0.036) | 1.515 (0.161) | 0.734 (0.012) | 0.967 (0.039) |

### Benchmarking AmpliDiff amplicons versus WGS

We show the effectiveness of amplicons selected by AmpliDiff by comparing abundance estimation results for SARS-CoV-2 lineages based on AmpliDiff amplicons, to abundance estimation results based on whole genome sequencing. To this end, we have generated two datasets: one for Texas and one for the Netherlands. Both datasets consist of a *reference set* (different from the reference set used for AmpliDiff) which contains the genomes that the abundance estimator uses as reference material, and a *simulation set* that contains genomes to simulate reads from. For both datasets, the reference genomes consist of genomes available on GISAID for the corresponding location, sampled between May and October of 2022. The simulation sets consist of genomes from the corresponding location with a sampling date in November 2022. A more detailed explanation of the setup for the simulation study can be found in the Methods section.

We perform abundance estimation using VLQ [4], a pipeline for estimating the relative abundance of viral lineages from wastewater sequencing data. VLQ employs kallisto [7] to assign a relative abundance to every genome in a set of reference genomes, based on the likelihood that reads originate from that genome. The relative abundance of an entire lineage is then obtained by aggregating the relative abundances of all genomes that are part of the lineage.

For both locations, we simulated sequencing data for amplicon widths of 200 and 400 bp, using the top 1, 2, 5, and 10 amplicons, or the whole genome. For each setting, we repeated the read simulation 20 times with different random seeds. For the amplicons selected by AmpliDiff, we check whether the corresponding primers bind (i.e. match exactly) to the input genomes—any sequences that do not have matching primers are not amplified, so we do not simulate reads in this case (see Methods). It is challenging, however, to maintain a similar approach for the whole genome-based simulations. There, we assume that both read coverage and read depth are uniform over the simulation genomes, and we do not check for primer binding. From a practical point of view, this is an ideal but highly unrealistic situation, particularly if we consider the context of wastewater sequencing [4]. Hence, the whole genome-based results should be considered as idealized results only serving as a benchmark for the amplicon-based abundance estimations.

Figure 3 shows the *Mean Absolute Prediction Errors* (MAPEs) which is used to evaluate abundance estimation accuracy for all datasets. We observe that the abundance estimations show very little variance, indicating that results are consistent in all configurations. In general, we see that the MAPEs range between 0.6% and 1.3% with the whole genome-based approach usually achieving the smallest MAPE. The exception to the latter observation is the Texas dataset using 5 or 10 amplicons of width 400 bp, or 10 amplicons of width 200 bp. In these cases, we see that the amplicon-based approach outperforms the whole genome-based approach, albeit by a small margin.

**Figure 3:**
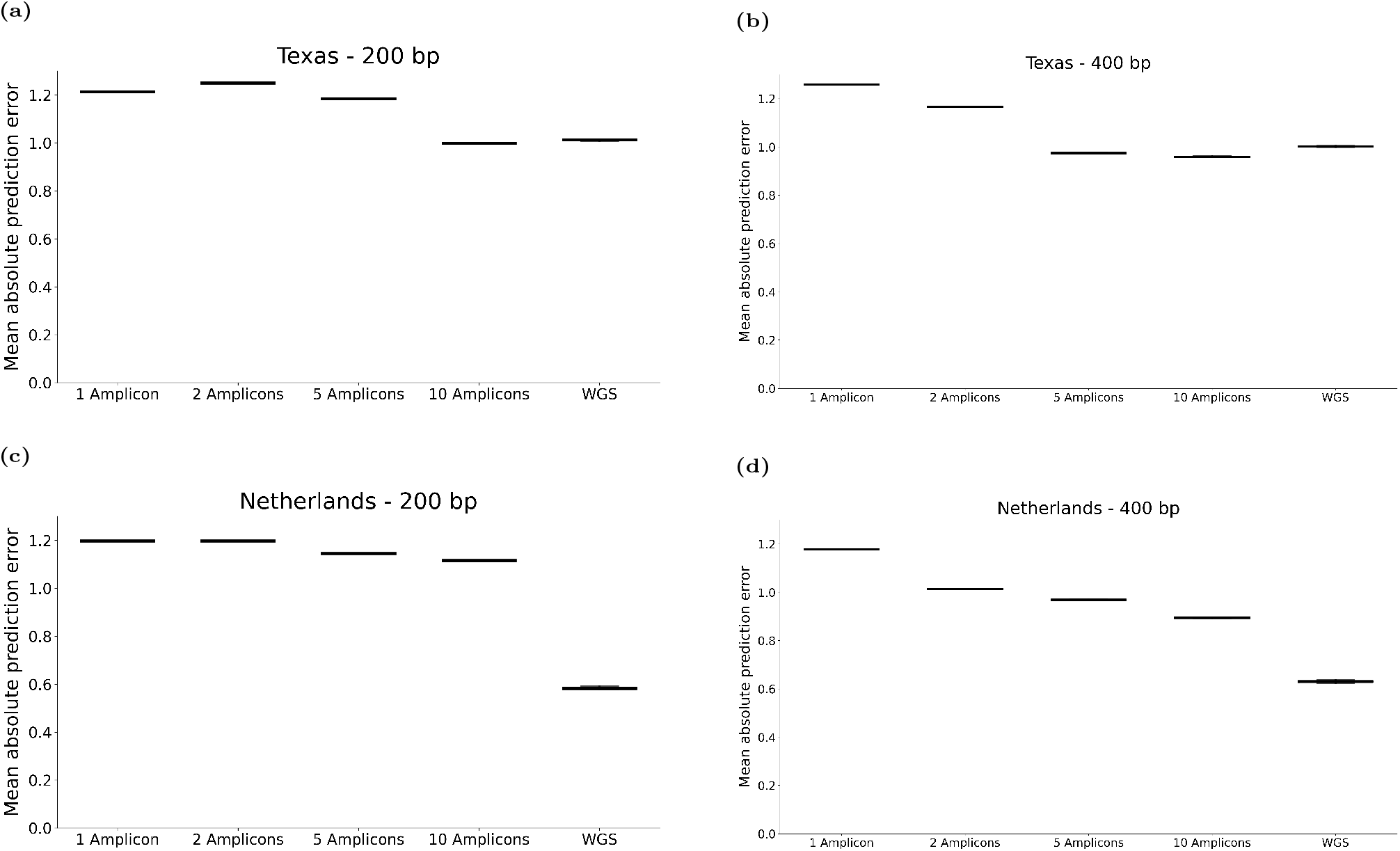
Mean absolute prediction errors for abundance estimations in both datasets for both amplicon widths using varying numbers of amplicons. **(a)** Texas dataset using amplicons of width 200. **(b)** Texas dataset using amplicons of width 400. **(c)** Netherlands dataset using amplicons of width 200. **(d)** Netherlands dataset using amplicons of width 400.

Interestingly, we see that both 200 and 400 bp amplicons yield similar results in the Texas dataset, whereas the 400 bp amplicons strongly outperform the 200 bp amplicons in the Netherlands dataset. In both regions we also see that using two amplicons of width 200 bp yields slightly worse MAPEs than using a single amplicon of width 200 bp, indicating that the second amplicon does not provide more information; instead, it seems to interfere with VLQ’s ability to correctly estimate relative abundances. Overall, we do see that adding amplicons reduces the MAPE.

In contrast to the Texas dataset, we see a substantial discrepancy between estimation results in the Netherlands dataset. This is particularly prominent for the amplicons of width 200 bp where, even with 10 amplicons, the whole genome-based approach strongly outperforms the amplicon-based approach. This can potentially be explained by the limited discriminatory power of short amplicons, or because of primers not binding. Alternatively, this discrepancy can be due to the fact that nearly all of the genomes used in the simulation study are (sub)lineages or recombinant lineages of the B.1.1.529 lineage (Omicron). Omicron has, since its first documentation in November 2021, rapidly become the dominant variant nearly everywhere [36], which explains its prominence in these experiments. Omicron sublineages can be particularly hard to distinguish in metagenomic samples as they are closely related and thus highly similar. Moreover, as most sublineages of B.1.1.529 in the simulation datasets did not exist in the PANGO nomenclature when the global reference set was constructed, the AmpliDiff amplicons were not explicitly designed to differentiate between these sublineages.

To see whether this is actually the case, we evaluate the prediction errors for individual lineages (Figures 4). Here, we consider each of the non-recombinant direct sublineages of B.1.1.529 as a single lineage (i.e. any lineage starting with B.1.1.529.1 is assigned to B.1.1.529.1). Lineages that do not occur in the reference set are not shown, since for those lineages all approaches have the same prediction errors. Figure4 shows that, at this resolution, most errors are due to wrongly estimating the abundances of B.1.1.529 sublineages. In addition, we see that the amplicon-based approach using 10 amplicons performs on par with the whole genome-based approach. This suggests that, if the main objective is to distinguish direct sublineages of B.1.1.529 from other lineages, the 10 amplicons found by AmpliDiff are able to do so. This observation is further supported by the results in Table 1, which shows the MAPE at this resolution for all approaches in both the Texas and Netherlands datasets. The amplicon-based approach using 10 amplicons outperforms the whole genome-based approach in all cases, which provides further evidence that the amplicon-based results are affected by the Omicron sublineages. We conclude that the amplicons selected by AmpliDiff can discriminate reasonably well between sublineages of B.1.1.529, and outperform the idealized whole genome-based approach on the resolution of discriminating between direct descendants of B.1.1.529.

**Figure 4:**
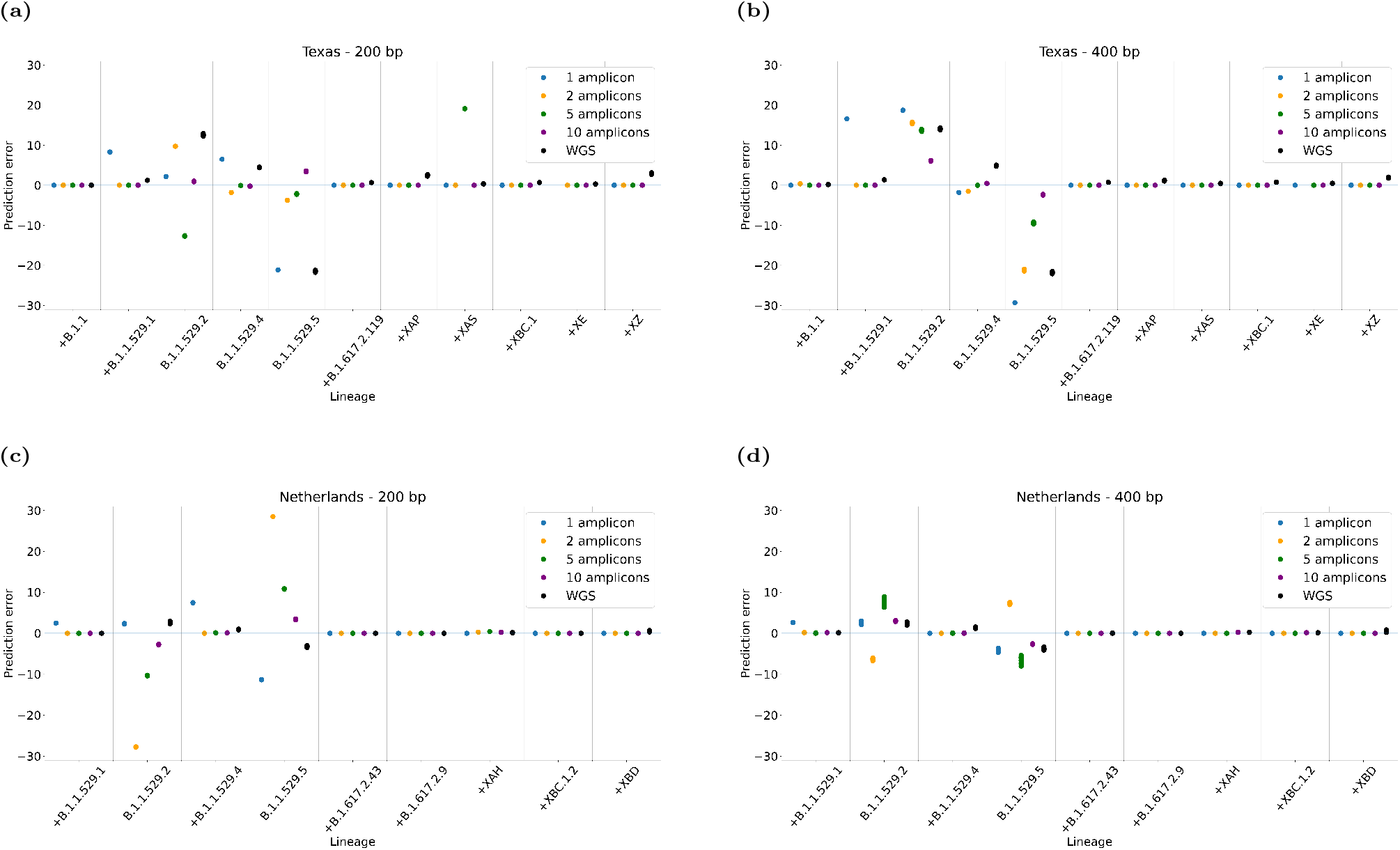
Prediction errors in % for both the amplicon-based approach (1, 2, 5, 10 amplicons) and the whole genome-based approach using sequencing fragments of lengths 200 and 400 in both datasets. The x-axes show the lineages (cut-off at most 3 sublineage levels) where a lineage prefixed by a ‘+’ character indicates a lineage that was not present in the simulation sample, and the y-axis is the prediction error.

### AmpliDiff is robust against missing data

The selection of reference genomes for metagenomics analyses is a common computational challenge. Due to the exponential growth of available genomes in genome databases [8], it can be necessary to concentrate only on a subset of genomes. On the other hand, there are also situations, such as the COVID-19 pandemic, where available information is limited and it can be impossible to find a reference set that is representative of the evolving population. In both scenarios, it is important to select an appropriate set of references that captures the diversity in the population, while not being excessively large (leading to inflated computational requirements). Consequently, when designing reference sets it is possible that informative reference genomes are left out, either due to limitations in computational resources or because of scarcity in the available information.

Here we consider what happens when AmpliDiff is presented with an incomplete reference set and the effect it has on the amplicons selected by AmpliDiff. For this, we take 15 random subsamples of sizes 1500 (54.6%), 2000 (72.8%), and 2500 (90.9%) respectively from the reference set used here (2749 reference genomes) and run AmpliDiff with the same settings as before.

Figures 5**(a)** and **(b)** show how often a nucleotide position is covered by an amplicon in a subsample, relative to the total number of subsamples, for amplicons of 200 bp and 400 bp width, respectively. As a general observation we see that, regardless of the number of excluded reference sequences and amplicon length, the majority of the results agree on which areas of the genome to cover. Moreover, the regions that are covered in at least 80% of these experiments also appear to overlap with the initial amplicons selected based on the full reference set. The variation in covered nucleotide positions that can be observed, however, stems from amplicons selected in the later stages of the greedy cycle. These amplicons generally contribute only marginally to the total differentiability of the selected amplicons and there are often multiple candidate amplicons with similar differentiability. Thus, the amplicon selection becomes ambiguous and therefore less consistent. Nevertheless, these results indicate that AmpliDiff is robust against missing reference genomes, even when nearly half of the reference genomes are excluded.

**Figure 5:**
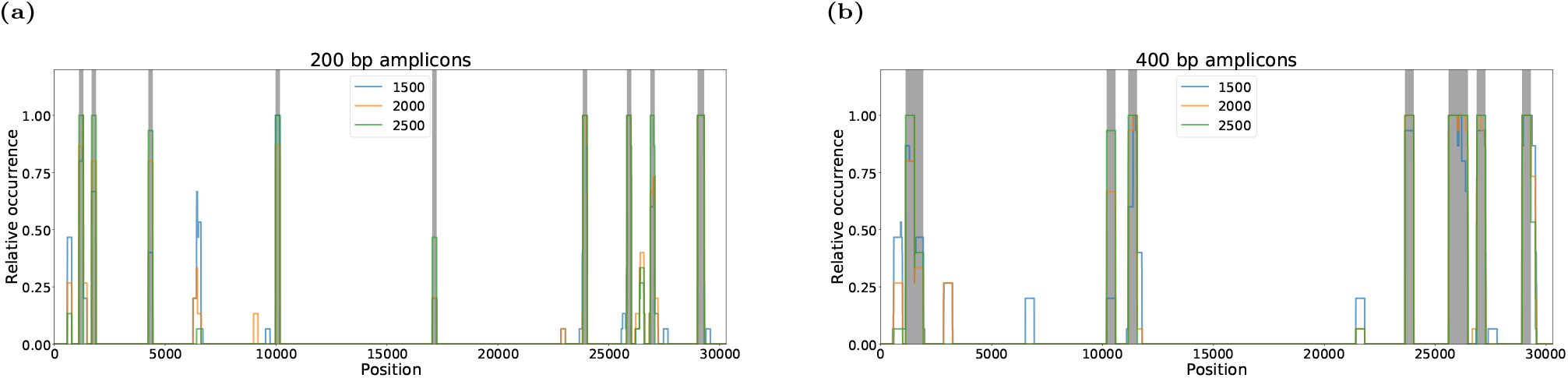
Assessment of the robustness of amplicons selected by AmpliDiff through subsampling subsets of sizes 1500, 2000, 2500, of the reference genomes 15 times for both amplicon widths. **(a)** shows for every position (in the aligned genomes) how often it was contained in an amplicon of width 200 in each of the subsampling experiments with grey regions indicating the amplicons on the full reference set, whereas **(b)** shows the same results for amplicons of width 400.

### Primers and amplicons found by AmpliDiff remain viable over time

A crucial aspect of amplicon design for the sake of discriminating between genomes, is that the primers also bind to genomes that are not present in the dataset used to determine the amplicons. We can check how well the amplicons determined by AmpliDiff work on ‘new’ genomes, by checking for every amplicon whether both a forward and reverse primer of the corresponding primer set bind. As a source of new genomes, we have downloaded high-quality genomes from GISAID from September 2022 to May 2023 (while the AmpliDiff input genomes were sampled up to August 2022) and checked the amplifiability of every amplicon in these genomes.

The amplifiability per month of every amplicon based on the 95% minimal amplifiablity requirement is shown in Figures 6 **(a)** and **(b)** for amplicon widths 200 and 400 bp, respectively. In Figure 6**(a)** we observe that every 200 bp amplicon retains an amplifiability of at least 95% in every month, with the exception of the final amplicon (17,042-17,242). This amplicon, located in ORF1b, appears to become significantly less amplifiable over time, reaching an amplifiability of approximately 70% in April 2023. Nevertheless, the amplifiability increases again in May 2023 and, as this is the final amplicon selected by AmpliDiff, it is expected not to have a significant effect on the ability to perform abundance estimation using the selected amplicons.

**Figure 6:**
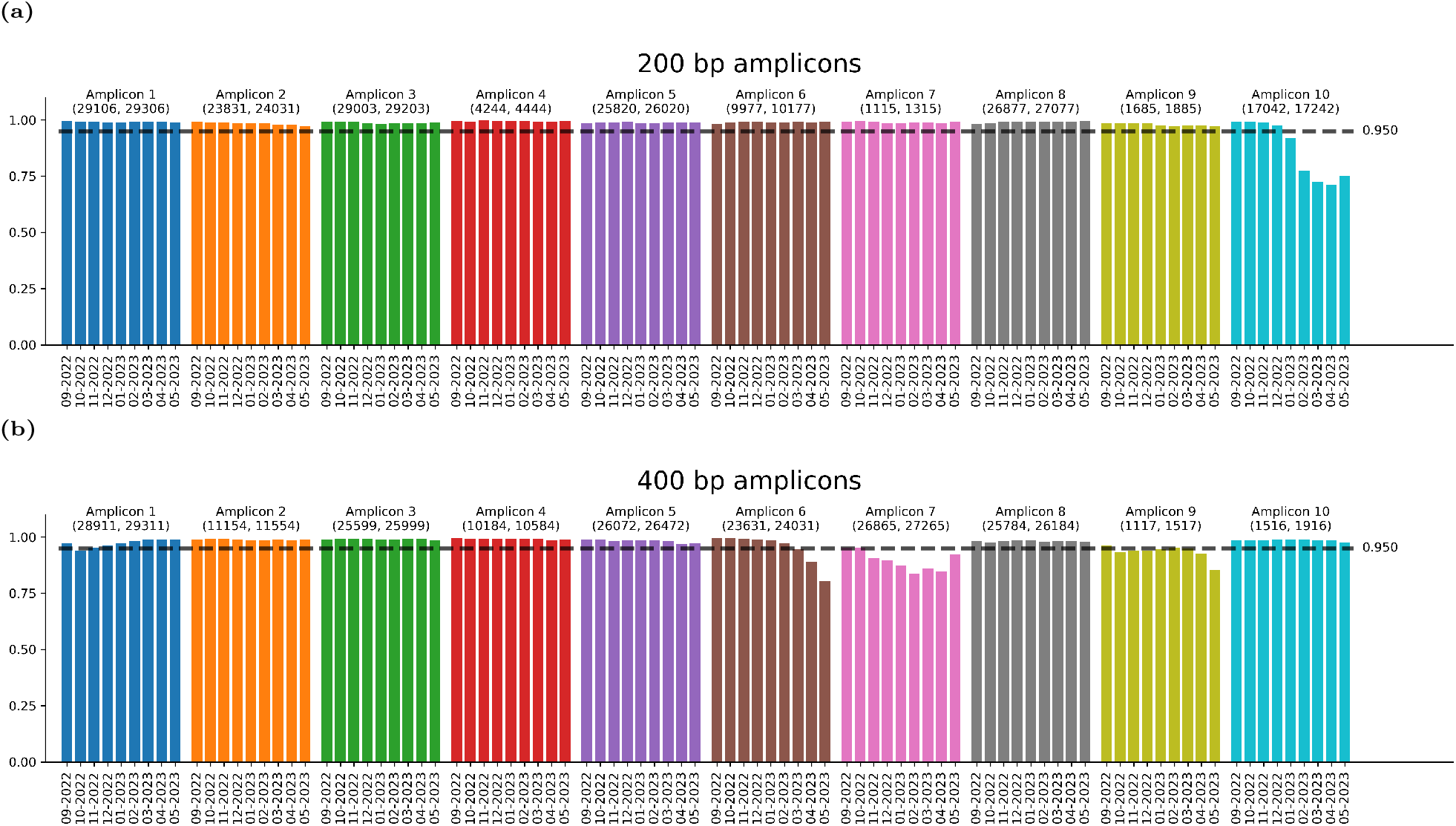
Fraction of genomes per month in which amplicons bind. **(a)** Amplifiability of 200-width amplicons generated with the 95% amplifiability requirement in the genomes from September 2022 to May 2023. **(b)** Amplifiability of 400 width amplicons generated with the 95% amplifiability requirement in the genomes from September 2022 to May 2023.

For amplicons of width 400 bp (Figure 6**(b)**) we see more variation in amplifiability, particularly in amplicons 6, 7, and 9. The pattern in amplifiability is also of interest, with amplicons 6 and 9 becoming less amplifiable over time, whereas amplicon 7 regains some of its amplifiability in May 2023 (similar to amplicon 10 in the 200 bp amplicons). As this again affects the later, less differentiable amplicons, we also expect that this does not strongly affect the abundance estimation accuracy using these amplicons.

Amplifiability results for amplicons based on different minimal amplifiability requirements are shown in Figures 7-12 in the Supplementary Material. We see similar trends in amplifiability, which is due to the fact that largely the same amplicons are chosen for different minimal required amplifiabilities. The exception to this are amplicons selected when the amplifiability is required to be 99.9% or 100%. In these cases, we observe that a single amplicon (although different between configurations) tends to be less amplifiable in both amplicon widths. Together, these observations indicate that the amplicons and primers found by AmpliDiff remain usable over time, with only a few relatively weakly differentiable amplicons experiencing some dropout.

### Computational requirements

All the results in this paper were obtained by running AmpliDiff on a High-Performance Computing (HPC) cluster, using 200 GB of RAM and 12 CPUs. Table 2 shows the average running time (over 15 runs) of AmpliDiff on 1500, 2000, and 2500 genomes for a minimal required amplifiability of 95%. The runtime of every step tends to increase with the number of included reference sequences, which is a natural consequence of the fact that each of the steps goes over all of the reference genomes.

**Table 2:** Average running time of AmpliDiff on an HPC cluster using 200 GB RAM and 12 CPU cores for different numbers of included genomes. Standard deviations are shown in parentheses, with the exception of the full reference set running times for which we performed only a single run.

| Task | 200 bp width |  |  |  | 400 bp width |  |  |  |
| --- | --- | --- | --- | --- | --- | --- | --- | --- |
|  | 1500 | 2000 | 2500 | 2749* | 1500 | 2000 | 2500 | 2749* |
| Constructing primer database | 390s (119s) | 682s (292s) | 695s (179s) | 673s | 446s (131s) | 548s (118s) | 695s (178s) | 818s |
| Determining feasible amplicons | 100s (22s) | 123s (23s) | 164s (42s) | 144s | 96s (22s) | 123s (29s) | 163s (35s) | 142s |
| Calculating amplicon differentiability | 151s (36s) | 274s (30s) | 440s (102s) | 526s | 194s (33s) | 351s (50s) | 560s (127s) | 602s |
| Greedy algorithm | 10,825s (2,878s) | 16,089s (3,062s) | 23,239s (4,926s) | 23,651s | 8,941s (2,376s) | 13,358s (3,085s) | 22,121s (4,134s) | 27,014s |

If we assume that determining the feasibility of a primer of length *k* takes *O*(*f* (*k*)) time, the primer database construction needs to parse every *k*-mer (*k* being equal to the primer length) in each of the reference genomes resulting in a theoretical runtime of *O*(|𝒢| · |*g*^*\**^| · *f* (*k*)), where | 𝒢 | is the number of included reference genomes and |*g*^*\**^| is the length of the largest genome. This is, however, a highly inflated runtime since the primer database keeps track of every *k*-mer that occurs by storing them in a dictionary. As a consequence, every subsequent time a *k*-mer occurs, it does not have to be checked for all its properties, thus reducing the runtime in practice.

Determining the feasible amplicons is done by considering every window of the amplicon size (either 200 or 400 bp here), and calculating the number of misalignment characters. This can be achieved in 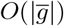 time for every multiple aligned genome 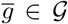 as every character has to be considered exactly once. Calculating the cumulative number of misalignment characters can also be done in 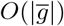 per genome which results in a theoretical runtime of 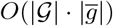 for determining the feasibility of amplicons. In a similar fashion, calculating the amplicon differentiability is done by moving a window with a size equal to the amplicon length over all pairs of genomes of different class and calculating the number of positions where a pair disagree, which yields a runtime of 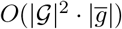. This term would seem to dominate the runtime of constructing the primer database, but we see from Table 2 that this is not the case in practice. While this may seem counter-intuitive, it is a consequence of the factor *f* (*k*) in the runtime of the primer database construction which accounts for checking all of the primer attributes, such as melting temperature, GC-content, and self-dimerization risk among others.

Finally, we should comment on the runtimes for the greedy algorithm. In contrast to the greedy algorithm for the set cover problem, it is difficult to provide a theoretical expected runtime for the greedy amplicon picking algorithm. This is due to two factors. First, we do not know which amplicons will have appropriate flanking primers a priori, which would require one to make assumptions about the likelihood of this being the case in order to come up with theoretical runtimes. The second factor ties into this, as AmpliDiff determines whether an amplicon can be amplified in enough reference genomes by solving a mixed integer linear program (MILP). While we do not have results on the hardness of the corresponding MILP, its relation to the set cover problem and maximum coverage problem suggests that it is likely NP-hard. Consequently, the runtime of deciding whether an amplicon can be added to the solution, and minimizing the number of primer pairs used for this relies on solving the MILP through a generic solver (Gurobi [17]), which we can assume to have an exponential runtime in the number of variables and/or constraints in the model. This explains why, in Table 2, we see that the greedy algorithm strongly dominates the overall runtime, with runtimes seemingly increasing exponentially with the number of included genomes.

## Discussion

Here, we introduce AmpliDiff, a computational tool designed to find discriminatory genomic regions along with primers, with the objective of discriminating between different lineages (or classes) of genomes. Without requiring prior knowledge of the input genomes beyond their respective lineages, AmpliDiff avoids relying on pre-defined regions of interest such as the 16S rRNA gene in bacteria and archaea [61], or the internal transcribed spacer (ITS) region in fungi [49]; instead, AmpliDiff can find combinations of genomic regions that collectively allow for the differentiation between genomes. By focusing only on discriminatory genomic regions, AmpliDiff provides an effective and economic alternative to amplification-based whole genome sequencing in the context of metagenomic profiling.

Through a simulation study, we show that the amplicons found by AmpliDiff can be used to obtain accurate and consistent abundance estimates for SARS-CoV-2 genomes. Moreover, we find that abundance estimation using AmpliDiff amplicons can generally achieve similar accuracy as abundance estimation based on whole genome sequencing, despite only using up to 10 amplicons. This is a major improvement over whole genome-based amplification for SARS-CoV-2, which requires approximately 100 amplicons and hence at least as many primer pairs.

Although AmpliDiff is based on a reference database, we see that amplicons selected by AmpliDiff remain highly discriminatory, even when the reference set used to determine the amplicons is not up to date. In particular, we observe that lineages get redefined over time, leading to misleading input for AmpliDiff: the amplicon selection approach may fail to distinguish between sequences from different lineages, as these were initially assigned to the same lineage. Hence, results could improve even further if the lineage assignment at the time of amplicon selection is stable.

A limitation of our benchmarking study is that we consider an idealized scenario for whole genome amplification, with complete genome coverage and uniform sequencing depth. While the simulated data for AmpliDiff amplicons takes the possibility of primers not binding (and hence amplicons not being amplified) into account, the simulated whole genome data does not. Achieving uniform coverage through amplicon-based sequencing is difficult to realize [2], which means that the WGS results are highly unrealistic to obtain in practice. This could be addressed by considering a similar setup where primer binding is taken into account during read simulation. The ARTIC primers have been widely used in practice and are regularly updated to account for new mutations occurring in the SARS-CoV-2 genome [35]. However, in our experiments, we check for exact primer binding, which the ARTIC primers are not designed for, and under our conditions, the ARTIC v4.1 amplicons would not amplify. If we were to incorporate less stringent primer binding conditions, we would expect to see two effects: 1) AmpliDiff amplicons may amplify better, likely improving results, and 2) the WGS-based results would likely suffer from issues such as amplicon dropout, which is expected to result in decreased performance for the WGS-based approach.

It is also of particular importance to experimentally show that primers found through primer design tools, AmpliDiff included, work in practice. This could be validated through in vitro experiments using synthetic populations and spiked-in samples of known variant composition. Such an approach would also lend itself to evaluating abundance estimation accuracy using AmpliDiff amplicons in practice. Moreover, in vitro experiments would allow for a fair comparison against abundance estimation based on ARTIC primers for example, and to verify the effectiveness of primers and amplicons found by AmpliDiff in the context of SARS-CoV-2.

In addition, it would also be of interest to apply AmpliDiff to sequence databases for other viruses, such as Influenza, SARS, measles, polio, or HIV. A prerequisite for any such analysis is a database with sufficiently many high-quality genomes. By using AmpliDiff for different species, we will gain additional insights into AmpliDiff’s effectiveness and limitations, allowing for further improvements to our methodology.

Finally, there are algorithmic aspects in AmpliDiff that can be improved. For example, the input genomes are pre-processed using MSA; since this is an NP-hard problem, we use a heuristic which can lead to sub-optimal alignments which in turn can affect amplicon selection. Moreover, MSA is not well suited for genomes with extensive recombination or repetitiveness, hence AmpliDiff is expected to work less well for complex genomes. Alternatively, pangenome representations (e.g. variation graphs [16] or colored de Bruijn graphs [23]) could serve as a data structure for AmpliDiff to work with. Another area for improvement is the fixed-length primer selection. In the current implementation, allowing variable-length primers would lead to a combinatorial explosion of the number of possible primer pairs. Finally, our primer feasibility algorithm relies on solving a MILP, which is related to the set cover problem and is therefore expected to be NP-hard. To this end, it would be valuable to prove hardness results for the amplicon selection problem and simultaneously work on finding dedicated algorithms for solving it efficiently.

## Conclusions

With AmpliDiff we introduce a computational tool to find genomic regions along with primers that can be used in the context of metagenomic profiling. Through a simulation study on SARS-CoV-2, we have shown that using amplicons found by AmpliDiff gives accurate and consistent abundance estimates of SARS-CoV-2 lineages, competitive with results based on WGS. Thus, AmpliDiff provides an effective and economic alternative to whole genome-based metagenomic profiling.

## Methods

### AmpliDiff methodology

The input to AmpliDiff consists of a reference set of multiple aligned genomes *G*, an amplicon width *L*_*A*_, a primer width *k*, a primer search window length *L*_0_, a maximum misalignment character threshold *L*_∅_, a maximal mismatch tolerance *D* and a minimal amplifiability threshold *γ*. In addition, AmpliDiff also accepts a list of classes for every genome in 𝒢 in TSV format (otherwise AmpliDiff assumes that the objective is to discriminate between all pairs of input genomes), a trade-off parameter *β* that is used to model the trade-off between adding primer pairs and the additional sequences in which an amplicon can be amplified, and finally a set of physicochemical properties that the primers have to satisfy. As shown in Figure 1, AmpliDiff consists of three steps: (i) a pre-processing step that builds a database of feasible primers for the reference genomes, (ii) an amplicon pre-processing step that extracts feasible amplicons and calculates their differentiability, and (iii) a greedy amplicon selection procedure which iteratively finds the most differentiable amplicon and adds it to the solution if primers can be found to amplify the amplicon. Here we give a description of these steps.

### Step (i): constructing the feasible primer database

The objective of AmpliDiff is to find amplicons along with primers such that they can be amplified with PCR. For this, it is necessary to consider the potential primer candidates (i.e. *k*-mers) that can be used from the genomes in 𝒢. By parsing the unaligned reference genomes (which can be obtained by removing misalignment characters) AmpliDiff extracts all *k*-mers, which are filtered according to the criteria listed in the Supplementary Material (including GC-content ranges, melting temperature ranges, risk of hairpin formation, and risk of self-dimerization). These properties are likely to affect the effectiveness of primers [14] and are therefore commonly considered in state-of-the-art primer design tools, such as Primer3 [54], OpenPrimeR [34] and PriMux [22]. The properties can be directly estimated from the nucleotide sequences of the primers. To prevent unwanted byproducts during PCR, primers that do not meet these criteria, as well as primers that occur at multiple locations in a single genome, are deemed infeasible.

After extracting all of the primer candidates, AmpliDiff filters the infeasible primers from the primer database. This results in a set of feasible primers 𝒫, which can be partitioned into forward primers 𝒫_*F*_ and reverse primers 𝒫_*R*_. An important property of AmpliDiff is that it allows reference genomes to contain degenerate nucleotides, represented by their corresponding IUPAC code [25]. When a genome contains a degenerate *k*-mer, AmpliDiff will try to disambiguate it by considering every possible representation of the *k*-mer. This approach differs from conventional approaches, where degenerate nucleotides are either replaced by pseudo-random nucleotides or kept degenerate, which potentially results in degenerate primers. If a *k*-mer occurring in a genome is too degenerate (i.e. has too many representative non-degenerate *k*-mers), this step is omitted, and the *k*-mer is skipped. The maximal number of different representations of a single *k*-mer can be defined by the user, and the default setting is 1024 different representations per *k*-mer. This approach always results in a set of non-degenerate primers which, through post-processing, could be turned into degenerate primers if desired.

#### Step (ii): amplicon pre-processing

By requiring that the input genomes *𝒢* are multiple aligned, it is guaranteed that similar and dissimilar regions in the genomes are aligned. Another consequence of multiple alignments is that all of the genomes in *𝒢* have the same length, which we will denote *L*_*G*_. In what follows we will also adhere to the following notation. Given a genome *g* ∈ *𝒢*, we define *g*[*i*] as the nucleotide at position *i* in genome *g* (starting with index 0). Similarly, for natural numbers *i < j* ≤ *L*_*G*_, we let *g*[*i* : *j*] = *g*[*i, i* + 1, …, *j* − 1] denote the substring of *g* starting at position *i* and ending before position *j*. Using this notation and the fact that all genomes in *𝒢* have the same length, we can define an amplicon *A* as an interval [*i* : *i* + *L*_*A*_], where *L*_*A*_ is the length of the amplicon. To simplify notation, we will write *g*[*A*] := *g*[*i* : *i* + *L*_*A*_] to refer to the substring defined by the interval *A* = [*i* : *i* + *L*_*A*_], in genome *g*.

In order for multiple aligned genomes to have the same length, they are padded by gap characters (‘-’). A gap character indicates that a genome has a gap at a given position, relative to (some of) the genomes it is aligned against. However, multiple sequence alignment is an NP-hard problem under general conditions [26]. Hence, most multiple sequence alignment programs (e.g. MAFFT [29] or ClustalW [53]) use heuristics, which generally provide sub-optimal alignments in practice. These alignments can have additional gaps, resulting in discriminatory regions based on falsely identified differences.

With AmpliDiff we combat this problem in two ways. First, we include a threshold *L*_∅_ for the maximum number of gap characters allowed per amplicon in all of the reference genomes. For this, we define ∅(*g*[*i* : *j*]) := |*{l* ∈ [*i, i* + 1, …, *j* − 2, *j* − 1] : *g*[*l*] = ‘-’*}* as the number of gap characters in genome *g*, on the interval [*i* : *j*]. Then, by requiring that ∅(*g*[*A*]) ≤ *L*_∅_ *∀g* ∈ *𝒢* for any amplicon *A*, we enforce that any input genome can only contain up to *L*_∅_ gap characters in *A*. By selecting an appropriate value for *L*_∅_ (default is 10% of the amplicon length), we remove amplicons that are likely affected by an inflated number of gap characters due to the sub-optimal multiple sequence alignment. Additionally, this ensures that the final PCR fragments will have approximately the same length.

In addition to classifying amplicons as infeasible due to large gaps, we apply another filter on amplicons in order to make sure that space directly flanking amplicons is available to find primers in. This is achieved by requiring that every amplicon is preceded and directly followed by at least *L*_*s*_ non-gap characters in every input genome. Together with the previous condition, this yields a set of feasible amplicon candidates which we will denote 𝒜.

The second way in which we alleviate the effect of sub-optimal alignments is by incorporating a maximal mismatch tolerance *D* (default value is 0) that controls when AmpliDiff considers a pair of genomes to be differentiated by an amplicon. Given an amplicon *A* = [*i* : *i* + *L*_*A*_] and two genomes *g, g*^*′*^ ∈ *𝒢* such that *g* ≁ *g*^*′*^(here ∼ is used as the equivalence relation between classes of genomes in *𝒢*), AmpliDiff considers the genomes to be differentiated by *A* if and only if |{*j* ∈ {*i*, …, *i* + *L*_*A*_ 1 : *g*[*j*] − *g*^*′*^[*j*]} | > *D*. In other words, an amplicon can only differentiate between a pair of genomes if they disagree at at least *D* + 1 nucleotide positions. Importantly, AmpliDiff only considers two characters *g*[*i*] and *g*^*′*^[*i*] to disagree if there is no overlap in the respective representations of *g*[*i*] and *g*^*′*^[*i*] when taking degenerate characters into account. This serves as a countermeasure against incorrectly selecting amplicons based on uncertainty in the input genomes.

Once the set of feasible amplicons, *𝒜*, has been determined, AmpliDiff continues by calculating the discriminatory power of amplicon candidates. For this, we define *δ*(*A*; *D*) := {(*g, g*^*′*^) *⊆ 𝒢* : |{*j* ∈ {*i*, …, *i* + *L*_*A*_ − 1} : *g*[*j*] ≠ *g*^*′*^[*j*]*}*| *> D}* as the set of genomes that can be differentiated by amplicon *A* under maximal mismatch tolerance *D*. Then, the *differentiability* of *A* is defined as Δ(*A*; *D*) := |*δ*(*A*; *D*)|. In the final step of step (ii), AmpliDiff calculates *δ*(*A*; *D*) and Δ(*A*; *D*) for every amplicon *A* ∈ 𝒜, which will serve as the starting point for the greedy amplicon selection step.

#### Step (iii): greedy amplicon selection

In the final step, AmpliDiff finds a near minimal set of amplicons with corresponding primers, to differentiate between as many pairs of genomes with different classes. The methodology employed by AmpliDiff is based on casting the amplicon finding problem as the *Set Cover Problem* (SCP) when looking to differentiate between all pairs of genomes of different classes, or the *Maximum Coverage Problem* (MCP) when looking to find a fixed number of amplicons that differentiate between as many pairs of class disjoint genomes as possible. These coverage problems are NP-hard combinatorial optimization problems [27, 32] that have been well-studied in the field of Operations Research. Instances of these coverage problems are given by a set of elements *𝒰*, often called the universe, a collection of subsets of the universe: *𝒞 ⊆* 2^*𝒰*^ such that *∪*_*S*∈*𝒞*_*S* = *𝒰* and a positive rational cost function *c* : *𝒞 →* ℚ_++_ that assigns a cost to every set in *𝒞*. In the SCP the objective is to find a minimal cost sub-collection *C*^*\**^ ⊆ 𝒞 such that ∪_*S*∈*𝒞*_*\** = 𝒰 whereas in the MCP the objective is to find a sub-collection of at most a given size covering as many elements of 𝒰 as possible.

To see how these coverage problems relate to the problem at hand, we can define our universe 𝒰 as the set of all pairs of genomes of different classes: 𝒰:= {*g, g*^*′*^} ⊆ 𝒢: *g* ≁ _*C*_ *g*^*′*^}. Then, defining an amplicon as the set of genomes it can differentiate and assigning every amplicon a cost of 1, we directly obtain an instance of the SCP, or an instance of the MPC if a maximum number of allowed amplicons is also given. As a consequence, we can use algorithms that have specifically been developed for these types of problems [9, 12, 19, 20, 64], to solve our amplicon finding problem. However, in our case, we also need to consider whether an amplicon can be amplified in sufficiently many genomes. Specifically, we require that an amplicon *A* ∈ 𝒜 has both a corresponding forward and reverse primer in at least *γ*% of the input genomes. Since we do not know a priori which amplicons are able to satisfy this condition, this has to be determined for every amplicon. It is computationally infeasible to do this beforehand for all candidate amplicons because we also need to check primer interactions (see below), hence we resort to a greedy approach similar to the greedy algorithm for the SCP and MCP [12]. Our approach iteratively adds the amplicon with the largest number of uncovered elements to the solution and whenever an amplicon is considered, we determine its amplifiability to see if it exceeds the minimal amplifiability condition. Note that the greedy algorithm for the SCP and MCP has been shown to be an approximation algorithm with approximation factor 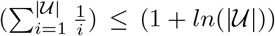, which means that if we were to determine which amplicons satisfy the minimal amplifiability constraint beforehand, we would obtain an approximately optimal solution for our problem as well.

The greedy amplicon selection step in AmpliDiff works as shown in Figure 1**(D)**. First, the candidate amplicons are sorted based on their differentiability (highest to lowest), and the most differentiable amplicon is picked, with ties broken arbitrarily. Using the primer database built in step (i), AmpliDiff finds feasible primers flanking this amplicon in all of the input genomes by solving a *Mixed Integer Linear Program* (MILP). The MILP is used to determine if it is possible to find primers that guarantee that the amplicon can be amplified in at least *γ*% of the input genomes (solving is done through Gurobi [17]). The MILP also checks whether it is possible to find enough forward and reverse primers that have approximately the same melting temperature (maximum difference of 5° Celsius) and do not pose a significant risk of having primer-primer interactions. Full details can be found in the Supplementary Material.

When an amplicon does not meet the minimal amplifiability requirement, it is discarded and the next most differentiable amplicon is considered. When an amplicon does meet the requirements, we solve the same MILP again, but now with the objective of finding a minimal set of primer pairs that enable the amplification of this amplicon in at least *γ*% of the genomes. If *γ* is smaller than 100%, we allow for an amplicon to not be amplifiable (i.e. have binding forward and reverse primers) in all of the reference genomes. Since the objective is to differentiate between pairs of genomes of different classes, we should in this case prioritize finding primers for genomes that contribute most to the differentiation power of the amplicon. Generic primer design tools do not consider this and thus we cannot rely on them to find primers.

To solve this problem we have added an auxiliary step in the greedy amplicon selection. In this step, we minimize the number of primer pairs required to amplify the candidate amplicon, while simultaneously maximizing the number of pairs of amplified genomes differentiated by the amplicon (see Supplementary Material). The trade-off between these conflicting objectives is modeled through the trade-off parameter *β* (default value 0.05) which allows adding additional primer pairs only if they increase the number of class disjoint genome pairs in which the amplicon can be amplified by at least 100 *· β*% of the total differentiability of the amplicon, as long as the amplifiability requirement is met. When the minimal required amplifiability is set to 100%, this trade-off is omitted since the amplicon is already required to be amplifiable in every input genome, and the number of required primer pairs is directly minimized instead.

After solving the primer minimization problem, the amplicon, along with the computed minimal set of primer pairs, is added to the solution. Then *δ*(*A*; *D*) is updated for the remaining amplicons in order to account for the pairs that can be differentiated by the added amplicon. Any amplicons with zero differentiability are removed and amplicons are again sorted based on their differentiability. The full amplicon selection loop continues until either all pairs of genomes with different classes can be differentiated, a given maximum number of amplicons have been found, or no more feasible amplicons exist, after which the found amplicons and their corresponding primer pairs are returned.

### Assessment on SARS-CoV-2

#### Amplicon generation

We have generated sets of 10 amplicons of different widths and with different minimal amplifiability requirements by using a global reference set of SARS-CoV-2 genomes. This reference set was constructed from all available genomes on the GISAID database [31] up until 18 August 2022, by following the reference set construction procedure detailed in [4]. This resulted in a set of 2,749 reference genomes distributed over 1,837 Pango lineages with 1-7 genomes per lineage. The reference genomes were multiple aligned using MAFFT [29] to obtain the input genomes for AmpliDiff. Then we ran AmpliDiff with *L*_*A*_ ∈ {200, 400}, *k* = 25, *L*_0_ = 50, *L*_∅_ = 0.1 *L*_*A*_, *D* = 1, *γ* ∈ {90%, 92.5%, 95%, 97.5%, 99.9%, 100%}, *β* ∈ {0.05, 0.10} and a maximum primer degeneracy of 1024 we obtained twelve sets of 10 amplicons of widths 200 and 400 bp and minimal amplifiabilities of 90%, 92.5%, 95%, 97.5%, 99.9%, and 100%, respectively (*β* did not affect results). The resulting amplicons can be found in Figures 1-6 in the Supplementary Material.

#### Simulation study

To evaluate how well amplicons and primers selected by AmpliDiff work in the context of lineage abundance estimation, we have created two benchmarking datasets: one with genomes from the Netherlands (constructed on 13 April 2023) and another with genomes from Texas (constructed on 6 March 2023). We estimate relative abundances of different lineages in these datasets using the amplicons selected by AmpliDiff, as well as for whole genome sequencing, by applying the VLQ pipeline [4]. Both datasets consist of a *reference dataset* with genomes spanning the months of May until October 2022 that is used by VLQ as references for the abundance estimations, and a *simulation dataset* with genomes from November 2022. The simulation dataset is used to simulate reads from where each simulation is repeated 20 times with different random seeds. This amounts to a total of 12 *×* 20 = 240 amplicon-based simulations and 2 *×* 20 = 40 whole genome-based simulations.

All datasets are filtered based on completeness and coverage to include only genomes with at least 29000 nucleotides which have at most 1% N-content and have fewer than 0.05% unique amino acid mutations (i.e. mutations not seen in other sequences in the GISAID database). The reference genomes were filtered further by following the VLQ pre-processing steps (see above), with the adjustment that they were not checked for ambiguous nucleotides.

For the whole genome-based abundance estimations, the reference genomes are directly used in VLQ. For the amplicon-based abundance estimation, we further process these reference genomes by extracting amplicons that are considered amplifiable in the respective genomes (see next subsection) and concatenating them with sequences of 200 ‘A’ nucleotides in between amplicons to avoid falsely mapping reads. This way, only the genomic regions corresponding to amplicons that we expect to bind are considered during the abundance estimation in VLQ.

Read simulations were done with ART [21], using the default mode for the whole genome-based reads and the amplicon mode for the amplicon-based reads. We have only simulated reads from amplicons that were amplifiable given the selected primers, whereas reads for the whole genome were simulated uniformly over the simulation genomes. For the fragments of length 200 (corresponding to 200 bp amplicons), we simulated 2 × 150 bp Illumina HiSeqX TrueSeq sequencing reads with an average fragment length of 200 bp and a standard deviation of 10 bp for the non-amplicon-based reads. Similarly, for fragments of length 400 (corresponding to 400 bp amplicons), we simulated 2 × 250 bp Illumina MiSeq v3 reads with an average fragment length of 400 bp and a standard deviation of 10 bp for the non-amplicon-based reads. For a fair comparison, we simulate datasets with a similar number of reads, hence we set the read depth to 1000× for the amplicon-based read simulations, and to 100× for the whole genome-based simulations. The lineage abundances in the simulated samples reflect the relative abundances of lineages in the simulation datasets.

We evaluate the abundance estimation accuracy by computing the *Mean Absolute Prediction Error*. For abundances {*ϕ*_l_}_*l*∈*ℒ*_ and estimated abundances 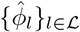, this is defined as 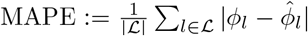 with ℒ equal to the set of all lineages occurring in the reference set or the simulation set.

### Primer binding and amplicon extraction

In our experiments, we assume that primers only bind to genomes if they match a genome exactly, as this is how primers are generated during step (i) of AmpliDiff. Since it is possible that primers bind to genomes in the presence of nucleotide mismatches in practice, this provides a conservative estimate of how amplifiable the amplicons found by AmpliDiff are in practice. However, when looking for nucleotide matches, we do allow up to 10 degenerate nucleotides in the primer binding site, which are considered to bind if their respective representations match the corresponding nucleotides in the primers. This approach may allow primers to bind to locations in the genome due to the presence of degenerate nucleotides, but we avoid discarding potential primer-binding sites. Note that the consequences are limited given that the input genomes are of high quality (i.e. low degeneracy).

## Declarations

### Ethics approval and consent to participate

Not applicable.

### Consent for publication

Not applicable.

### Availability of data and materials

The reference genomes used for determining the AmpliDiff amplicons are available through GISAID under identifier EPI SET 230825gp. The reference genomes and genomes used to simulate reads from for the Texas-based simulation experiments are available through GISAID under identifiers EPI SET 230825zm and EPI SET 230825pe, respectively. The reference genomes and genomes used to simulated reads from for the Netherlands-based simulation experiments are available through GISAID under identifiers EPI SET 230825he and EPI SET 230825fe, respectively. The simulated sequencing data (.fastq files) for the 95% amplifiability amplicons and whole genome sequencing approaches are available on Zenodo (https://zenodo.org/record/8298887) [56]. The code for AmpliDiff can be found on github (https://github.com/JaspervB-tud/AmpliDiff) under the MIT license [55].

### Authors’ contributions

J.A.B. and D.S.S. conceived the study. J.B. and J.A.B. developed the method with feedback from D.S.S. J.B. implemented the method and performed data analysis. J.B. and J.A.B. wrote the manuscript with feedback from D.S.S. All authors read and approved the final manuscript.

## Acknowledgements

We gratefully acknowledge all data contributors, i.e., the Authors and their Originating laboratories responsible for obtaining the specimens, and their submitting laboratories for generating the genetic sequence and metadata and sharing via the GISAID Initiative, on which this research is based.

## Supplementary Information

### Additional file 1

Includes all supplementary information, supplementary figures and supplementary tables.

