## Supplementary Information for "AmpliDiff: An Optimized Amplicon Sequencing Approach to Estimating Lineage Abundances in Viral Metagenomes"

### Cumulative differentiability plots

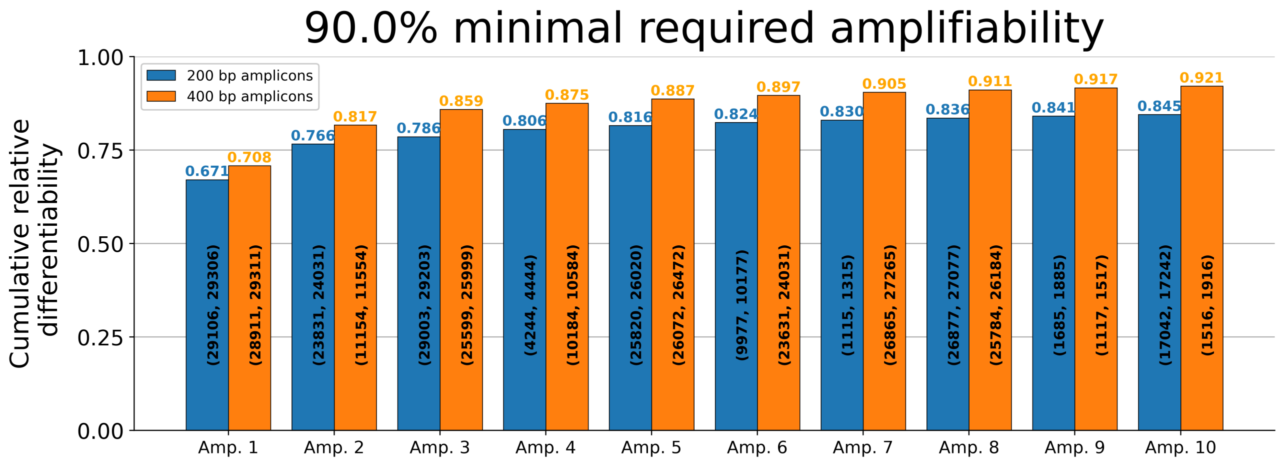


**Figure 1:** Cumulative relative differentiability of ten best amplicons (both 200 and 400 widths) found by AmpliDiff with the requirement of at least 90% amplifiability

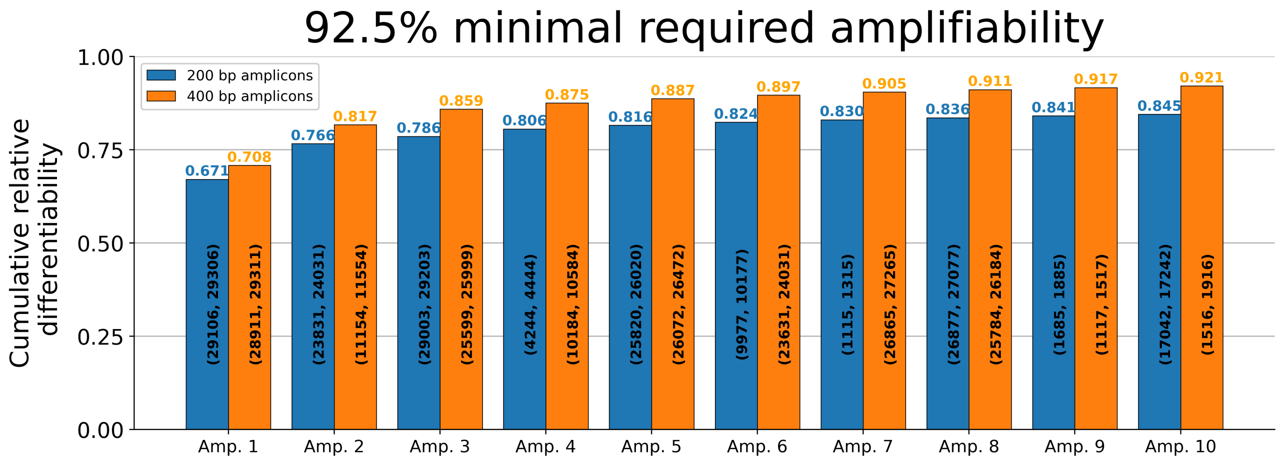


**Figure 2:** Cumulative relative differentiability of ten best amplicons (both 200 and 400 widths) found by AmpliDiff with the requirement of at least 92.5% amplifiability

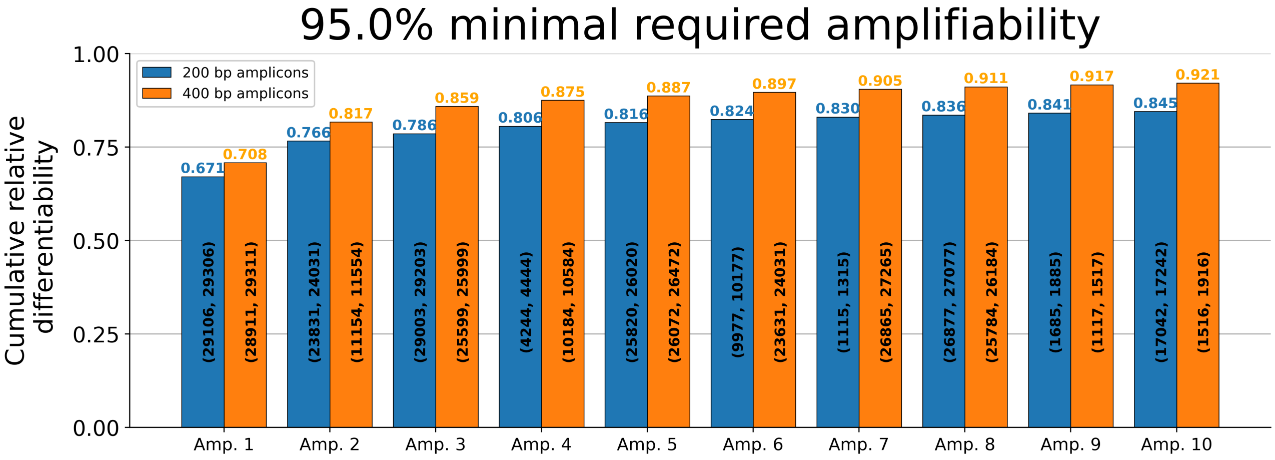


**Figure 3:** Cumulative relative differentiability of ten best amplicons (both 200 and 400 widths) found by AmpliDiff with the requirement of at least 95% amplifiability

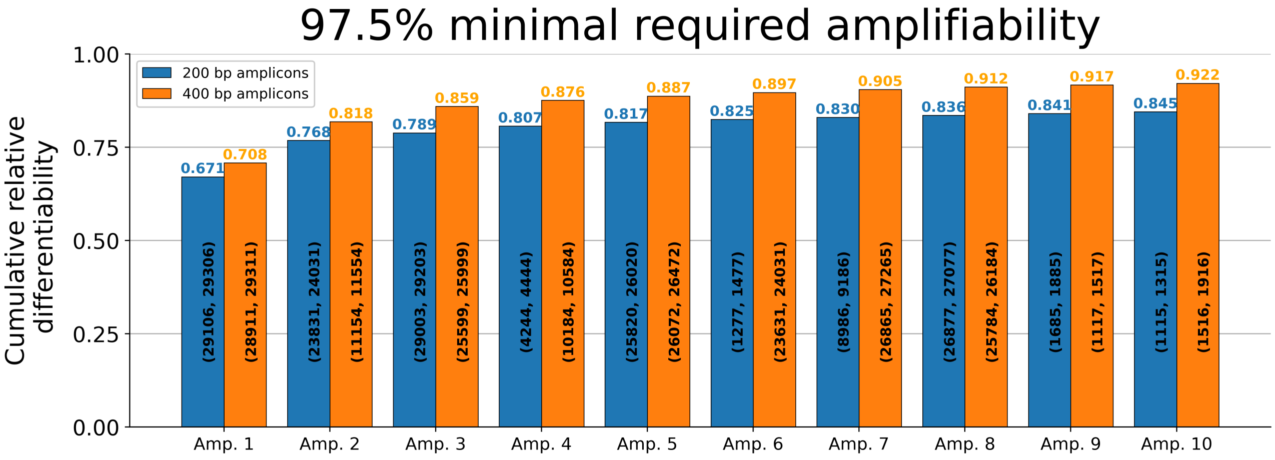


**Figure 4:** Cumulative relative differentiability of ten best amplicons (both 200 and 400 widths) found by AmpliDiff with the requirement of at least 97.5% amplifiability

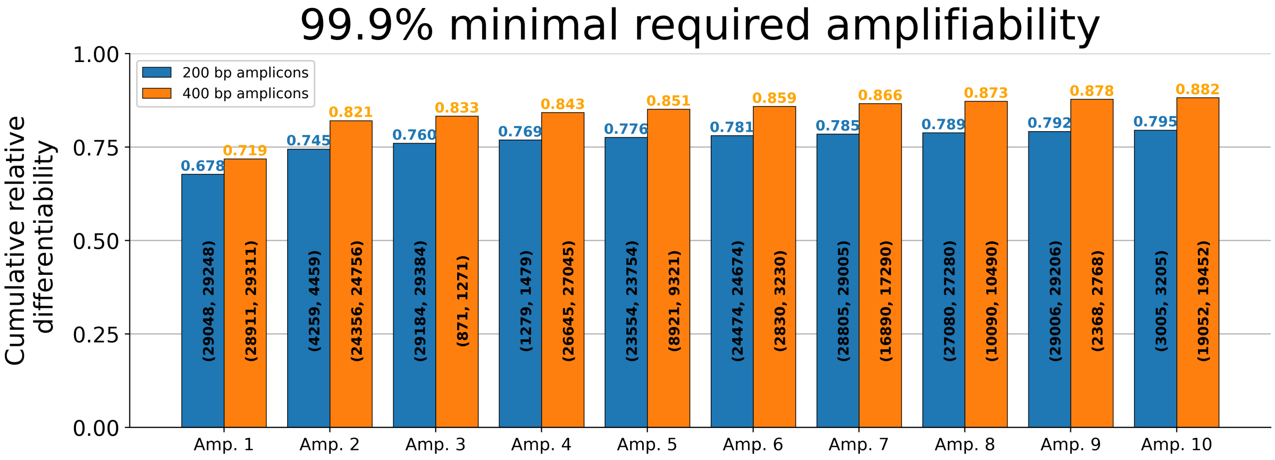


**Figure 5:** Cumulative relative differentiability of ten best amplicons (both 200 and 400 widths) found by AmpliDiff with the requirement of at least 99.9% amplifiability

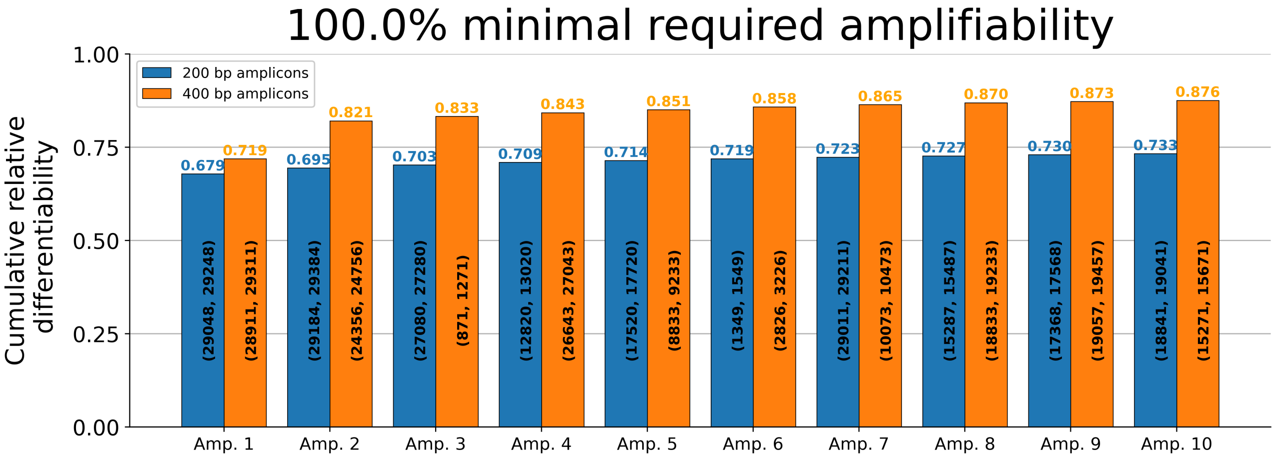


**Figure 6:** Cumulative relative differentiability of ten best amplicons (both 200 and 400 widths) found by AmpliDiff with the requirement of 100% amplifiability

**(a)**

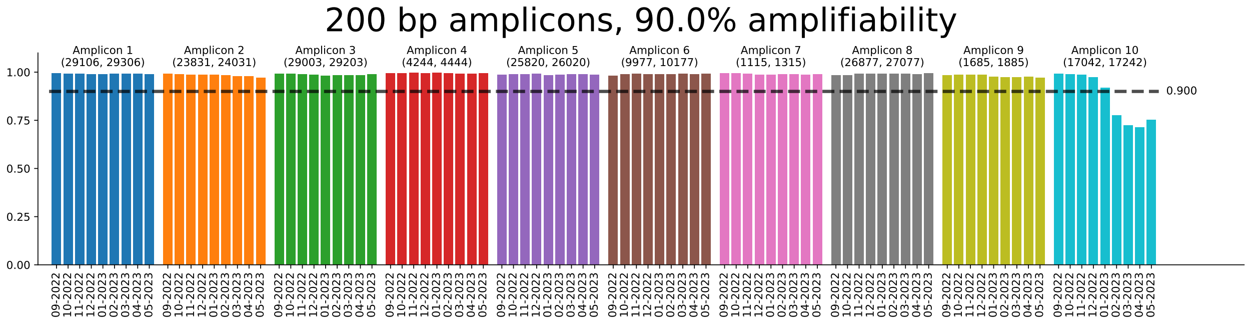


**(b)**

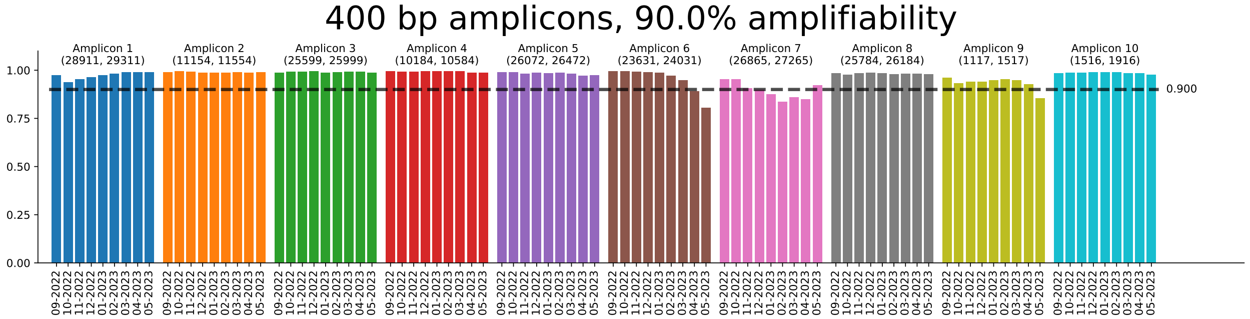


**Figure 7:** Fraction of genomes per month in which amplicons bind. **(a)** Amplifiability of 200 width amplicons generated with a 90% amplifiability requirement, in the "out of sample" genomes from September 2022 to May 2023. **(b)** Amplifiability of 400 width amplicons generated with a 90% amplifiability requirement, in the "out of sample" genomes from September 2022 to May 2023.

### Required primer pairs

| **Amplicon width** | **200** | | | | | | **400** | | | | | |
|---|---|---|---|---|---|---|---|---|---|---|---|---|
| **Minimal amplifiability** | 90% | 92.5% | 95% | 97.5% | 99.9% | 100% | 90% | 92.5% | 95% | 97.5% | 99.9% | 100% |
| Amplicon 1 | 1F, 1R | 1F, 1R | 1F, 1R | 1F, 1R | 7F, 6R | 9F, 6R | 1F, 1R | 1F, 1R | 1F, 1R | 1F, 1R | 3F, 4R | 3F, 6R |
| Amplicon 2 | 1F, 1R | 1F, 1R | 1F, 1R | 4F, 3R | 11F, 5R | 11F, 4R | 1F, 1R | 1F, 1R | 1F, 1R | 2F, 2R | 5F, 3R | 7F, 3R |
| Amplicon 3 | 2F, 2R | 2F, 2R | 2F, 2R | 3F, 3R | 9F, 4R | 7F, 5R | 1F, 1R | 1F, 1R | 1F, 1R | 1F, 1R | 5F, 5R | 6F, 7R |
| Amplicon 4 | 1F, 1R | 1F, 1R | 1F, 1R | 1F, 1R | 9F, 9R | 3F, 8R | 2F, 2R | 2F, 2R | 2F, 2R | 2F, 2R | 1F, 5R | 1F, 7R |
| Amplicon 5 | 1F, 1R | 1F, 1R | 1F, 1R | 2F, 2R | 4F, 6R | 13F, 4R | 1F, 1R | 1F, 1R | 1F, 1R | 1F, 1R | 5F, 5R | 5F, 9R |
| Amplicon 6 | 1F, 1R | 1F, 1R | 1F, 1R | 1F, 1R | 4F, 2R | 2F, 4R | 1F, 1R | 1F, 1R | 1F, 1R | 1F, 1R | 5F, 6R | 5F, 6R |
| Amplicon 7 | 1F, 1R | 1F, 1R | 1F, 1R | 1F, 1R | 2F, 9R | 4F, 7R | 2F, 2R | 2F, 2R | 2F, 2R | 2F, 2R | 4F, 2R | 3F, 9R |
| Amplicon 8 | 2F, 2R | 2F, 2R | 2F, 2R | 2F, 2R | 5F, 5R | 3F, 5R | 1F, 1R | 1F, 1R | 1F, 1R | 3F, 3R | 4F, 4R | 10F, 8R |
| Amplicon 9 | 1F, 1R | 1F, 1R | 1F, 1R | 2F, 2R | 3F, 5R | 5F, 7R | 1F, 1R | 1F, 1R | 1F, 1R | 1F, 1R | 3F, 7R | 9F, 3R |
| Amplicon 10 | 1F, 1R | 1F, 1R | 1F, 1R | 1F, 1R | 12F, 7R | 10F, 5R | 1F, 1R | 1F, 1R | 1F, 1R | 1F, 1R | 7F, 2R | 5F, 5R |

**Table 1:** Number of primers corresponding to every amplicon found by AmpliDiff for different amplicon widths and minimal required amplifiabilities. F denotes the number of forward primers and R is the number of reverse primers.

**(a)**

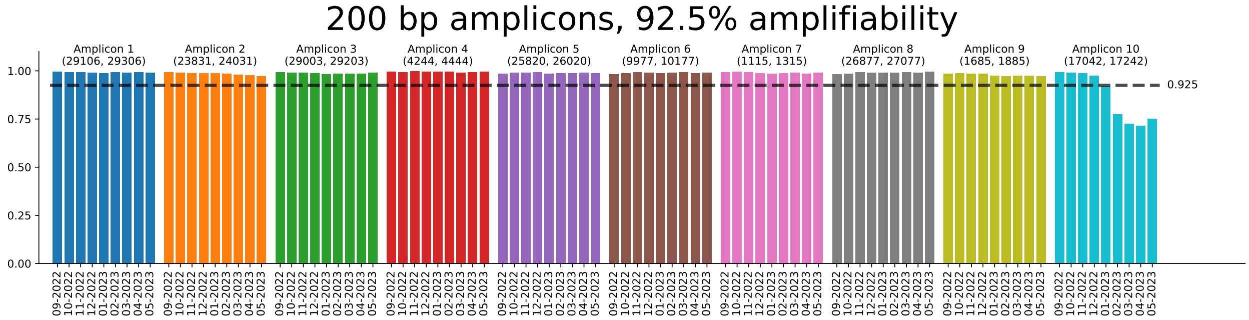


**(b)**

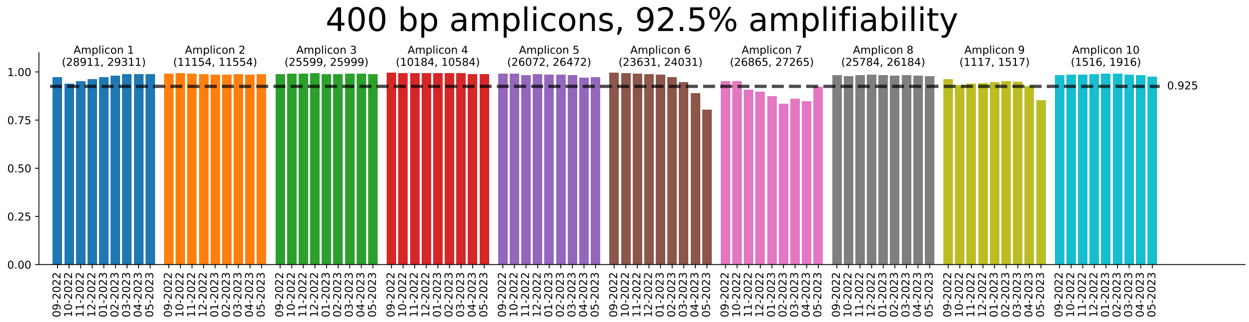


**Figure 8:** Fraction of genomes per month in which amplicons bind. **(a)** Amplifiability of 200 width amplicons generated with a 92.5% amplifiability requirement, in the "out of sample" genomes from September 2022 to May 2023. **(b)** Amplifiability of 400 width amplicons generated with a 92.5% amplifiability requirement, in the "out of sample" genomes from September 2022 to May 2023.

**(a)**

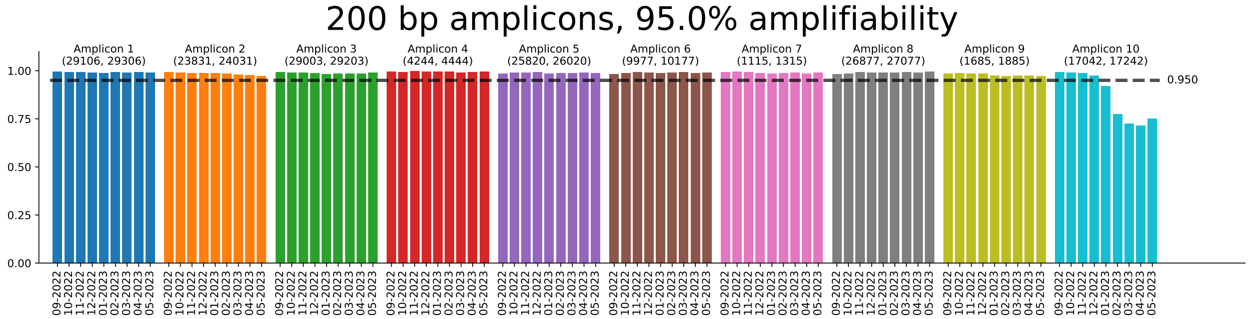


**(b)**

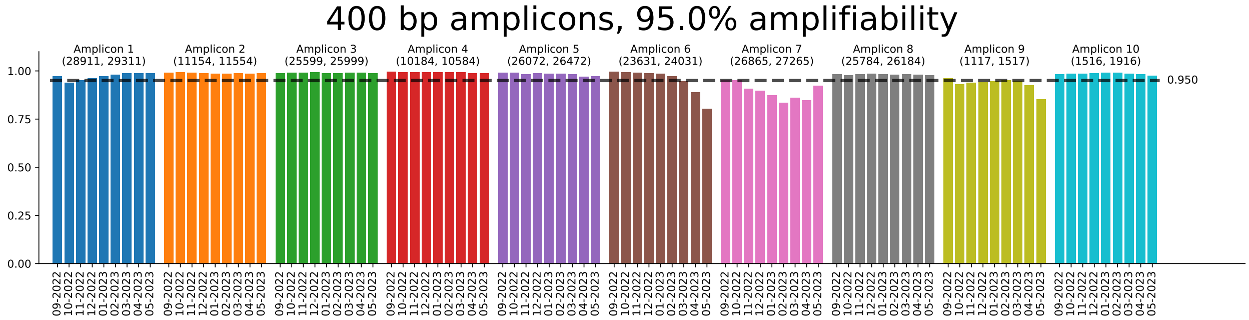


**Figure 9:** Fraction of genomes per month in which amplicons bind. **(a)** Amplifiability of 200 width amplicons generated with a 95% amplifiability requirement, in the "out of sample" genomes from September 2022 to May 2023. **(b)** Amplifiability of 400 width amplicons generated with a 95% amplifiability requirement, in the "out of sample" genomes from September 2022 to May 2023.

**(a)**

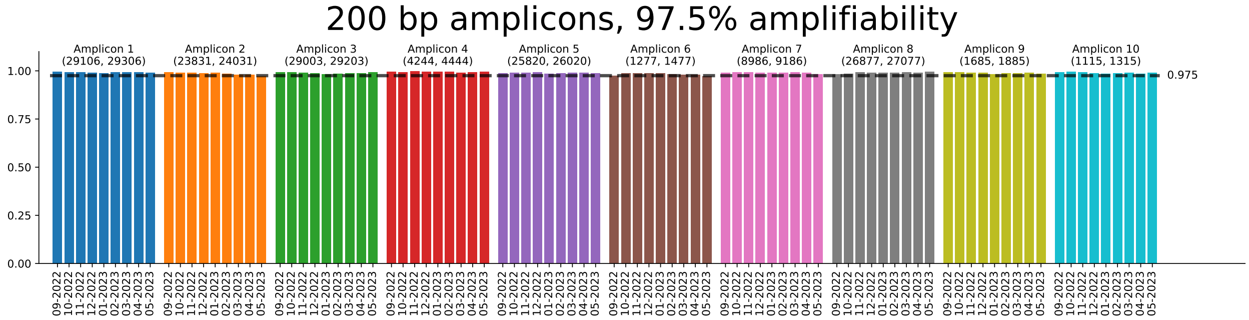


**(b)**

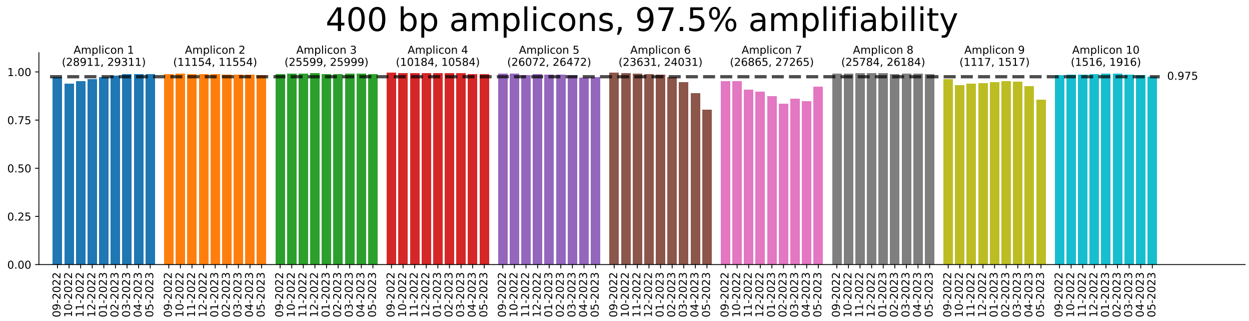


**Figure 10:** Fraction of genomes per month in which amplicons bind. **(a)** Amplifiability of 200 width amplicons generated with a 97.5% amplifiability requirement, in the "out of sample" genomes from September 2022 to May 2023. **(b)** Amplifiability of 400 width amplicons generated with a 97.5% amplifiability requirement, in the "out of sample" genomes from September 2022 to May 2023.

**(a)**

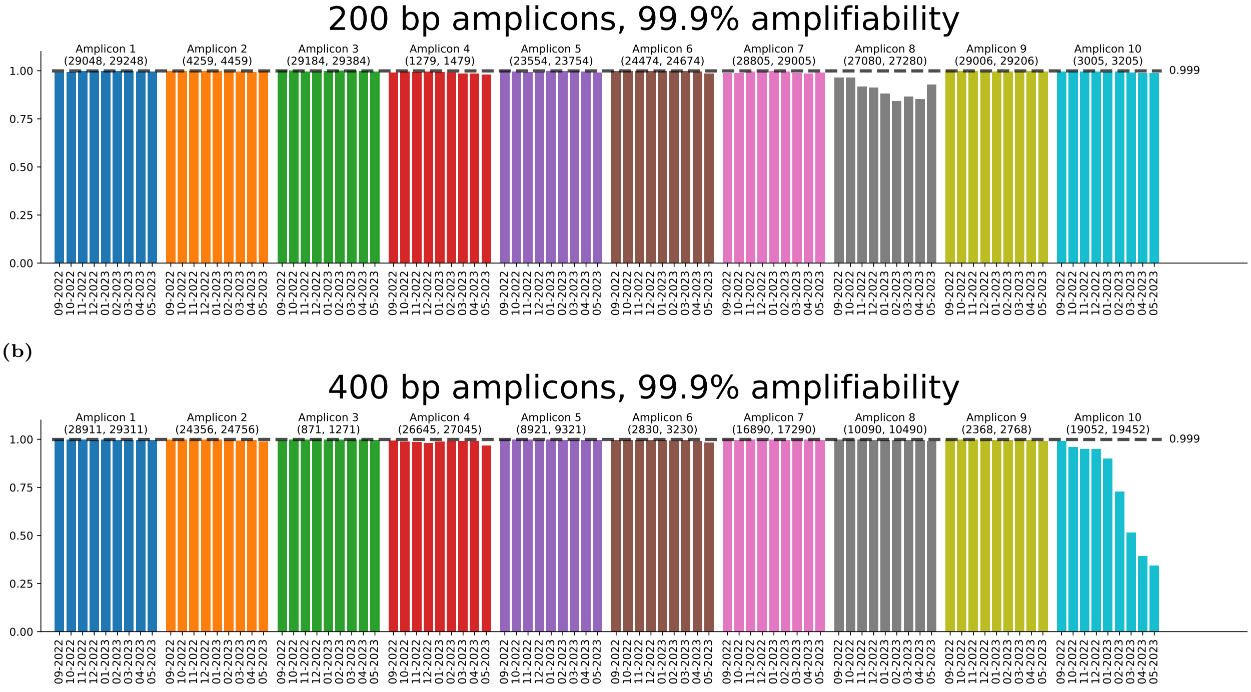


**Figure 11:** Fraction of genomes per month in which amplicons bind. **(a)** Amplifiability of 200 width amplicons generated with a 99.9% amplifiability requirement, in the "out of sample" genomes from September 2022 to May 2023. **(b)** Amplifiability of 400 width amplicons generated with a 99.9% amplifiability requirement, in the "out of sample" genomes from September 2022 to May 2023.

**(a)**

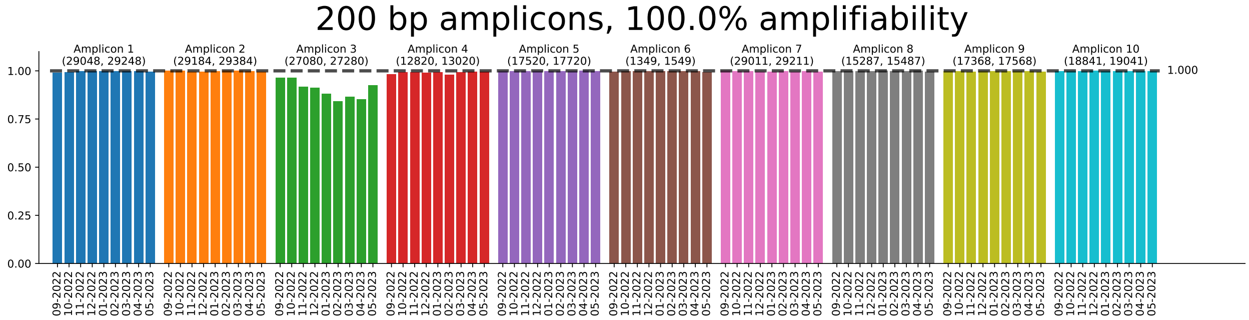


**(b)**

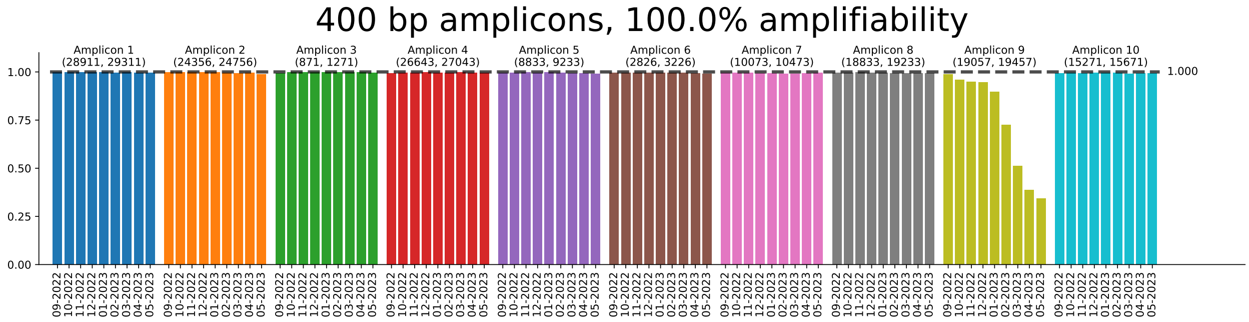


**Figure 12:** Fraction of genomes per month in which amplicons bind. **(a)** Amplifiability of 200 width amplicons generated with a 100% amplifiability requirement, in the "out of sample" genomes from September 2022 to May 2023. **(b)** Amplifiability of 400 width amplicons generated with a 100% amplifiability requirement, in the "out of sample" genomes from September 2022 to May 2023.

### Primer binding results

### Primer selection criteria

- GC-content in range 40-60%
- Melting temperature in the range of 55-75degC
- Maximum of 2 A/T characters in final 3 nucleotides (3'-end)
- Maximum of 3 G/C characters in final 5 nucleotides (3'-end)
- Maximum run of 3 (i.e. 3 consecutive identical nucleotides)
- Maximum run of 2 duo nucleotides (e.g. ACAC)
- MFE threshold -5 (a proxy for the risk of hairpin formation)
- At most 10 self-complementary basepairs when compared to its own reverse complement (this is for the "worst" alignment)

### Primer feasibility/optimization model

$$\max \sum_{(s,s')\in\mathcal{D}} y_{(s,s')} - \beta \cdot |\mathcal{D}| \cdot Q$$

$$\text{s.t.}$$

$$z_s \leq \sum_{p\in\mathcal{P}_s^f} x_p \qquad \forall s \in \mathcal{S}$$

$$z_s \leq \sum_{p\in\mathcal{P}_s^r} x_p \qquad \forall s \in \mathcal{S}$$

$$\sum_{s\in\mathcal{S}} z_s \geq \alpha|\mathcal{S}|$$

$$Q \geq \sum_{p\in\mathcal{P}^f} x_p$$

$$Q \geq \sum_{p\in\mathcal{P}^r} x_p$$

$$y_{(s,s')} \leq 0.5z_s + 0.5z_{s'} \qquad \forall (s,s') \in \mathcal{D}$$

$$T^+ \geq T_p x_p \qquad \forall p \in \mathcal{P}^f \cup \mathcal{P}^r$$

$$T^- \leq T_p(3 - 2x_p) \qquad \forall p \in \mathcal{P}^f \cup \mathcal{P}^r$$

$$T^+ - T^- \leq T^*$$

$$x_p + x_{p'} \leq C(p,p') \qquad \forall (p,p') \in (\mathcal{P}^f \cup \mathcal{P}^r) \times \big((\mathcal{P}^f \cup \mathcal{P}^r) \setminus p\big)$$

$$x_p \in \{0,1\} \forall p \in \mathcal{P}^f \cup \mathcal{P}^r$$

$$z_s \in \{0,1\} \forall s \in \mathcal{S}$$

$$T^+ \geq 0$$

$$T^- \geq 0$$

##### Model variables and parameters

#### Variables:

- $x_p$ : binary variable equal to 1 if primer $p \in \mathcal{P}$ is selected
- $z_s$ : binary variable equal to 1 if sequence $s$ has at least one binding forward and one binding reverse primer

- $y_{(s,s')}$ : binary variable equal to 1 if both sequences $s$ and $s'$ have at least one binding forward and one binding reverse primer
- $T^*$ : continuous variable equal to the difference between the highest and lowest melting temperature of chosen primers
- $Q$ : integer-valued variable equal to the number of primer pairs chosen

**Variable sets:**

- $\mathcal{D}$ : set of all pairs of sequences of different class/lineage that can be differentiated by the current amplicon
- $\mathcal{S}$ : set of all sequences
- $\mathcal{P}^f$ : set of all forward primers
- $\mathcal{P}^r$ : set of all reverse primers
- $\mathcal{P}^f_s$ : set of forward primers corresponding to sequence $s$
- $\mathcal{P}^r_s$ : set of all reverse primers corresponding to sequence $s$

**Parameters:**

- $\alpha$ : fraction of input genomes in which the amplicon must be amplifiable
- $\beta$ : trade-off parameter between the number of included primer pairs and additional differentiability
- $T^+$ : maximum allowed melting temperature
- $T^-$ : minimum allowed melting temperature
- $T_p$ : approximate melting temperature of primer $p$
- $C(p, p')$ : binary value equal to 1 if primers $p$ and $p'$ are predicted to form a primer-dimer pair

##### Constraints explanation

The first two sets of constraints set $z_s$ equal to 1 if there primers are selected to amplify the amplicon in $s$ (forward and reverse). The third constraint set enforces the amplicon amplifiability. Constraint sets 4 and 5 enforce that $Q$ is equal to the number of primer pairs (i.e. the maximum of the number of forward and reverse primers). Since the objective minimizes $Q$ it is only necessary to include these "greater or equal" constraints. Constraint set 6 enforces that $y_{(s,s')}$ is only set to 1 if both sequences $s$ and $s'$ satisfy amplifiability. Constraint sets 7 through

9 model the melting temperature constraints. Note that constraint set 8 uses $T^- \leq T_p(3-2x_p)$ which works as primer temperatures are generally in the range 40-70 and thus if $x_p$ is not selected the constraint is redundant, and if $x_p$ is selected then it sets $T^-$ to at most $T_p$. Finally, constraint set 10 enforces that incompatible primers can not simultaneously be added to the solution.

This version of the model considers the optimization of the number of primer pairs needed for amplification of an amplicon, taking into account in which sequences the amplicon can be amplified. Note that in the feasibility check, the objective value is omitted as it is only required to check if a feasible solution exists. When the value of $\alpha$ is equal to 1, the model is simplified by removing the variables $y_{(s,s')}$ (and corresponding constraints) as these will be forced to equal 1. Additionally, the objective in this case is simply to minimize the number of required primer pairs $Q$.
